# Global-scale CRISPR gene editor specificity profiling by ONE-seq identifies population-specific, variant off-target effects

**DOI:** 10.1101/2021.04.05.438458

**Authors:** Karl Petri, Daniel Y. Kim, Kanae E. Sasaki, Matthew C. Canver, Xiao Wang, Hina Shah, Hyunho Lee, Joy E. Horng, Kendell Clement, Sowmya Iyer, Sara P. Garcia, Jimmy A. Guo, Gregory A. Newby, Luca Pinello, David R. Liu, Martin J. Aryee, Kiran Musunuru, J. Keith Joung, Vikram Pattanayak

**Affiliations:** Molecular Pathology Unit, Massachusetts General Hospital, Charlestown, MA, USA.; Center for Cancer Research, Massachusetts General Hospital, Charlestown, MA, USA.; Center for Computational and Integrative Biology, Massachusetts General Hospital, Charlestown, MA, USA.; Department of Pathology, Harvard Medical School, Boston, MA, USA.; Merkin Institute of Transformative Technologies in Healthcare, Broad Institute of Harvard and MIT, Cambridge, MA, USA.; Department of Chemistry and Chemical Biology, Harvard University, Cambridge, MA, USA.; Howard Hughes Medical Institute, Harvard University, Cambridge, MA, USA.; Division of Cardiology and Cardiovascular Institute, Department of Medicine, Perelman School of Medicine at the University of Pennsylvania, Philadelphia, PA, USA.; Department of Biostatistics, Harvard T.H. Chan School of Public Health, Boston, MA, USA.

## Abstract

Defining off-target profiles of gene-editing nucleases and CRISPR base editors remains an important challenge for use of these technologies, therapeutic or otherwise. Existing methods can identify off-target sites induced by these gene editors on an individual genome but are not designed to account for the broad diversity of genomic sequence variation that exists within populations of humans or other organisms. Here we describe OligoNucleotide Enrichment and sequencing (**ONE-seq**), a novel *in vitro* method that leverages customizable, high-throughput DNA synthesis technology instead of purified genomic DNA (**gDNA**) from individual genomes to profile gene editor off-target sites. We show that ONE-seq matches or exceeds the sensitivity of existing single-genome methods for identifying *bona fide* CRISPR-Cas9 off-target sites in cultured human cells and *in vivo* in a liver-humanized mouse model. In addition, ONE-seq outperforms existing best-in-class single-genome methods for defining off-target sites of CRISPR-Cas12a nucleases, cytosine base editors (**CBE**s), and adenine base editors (**ABE**s), unveiling previously undescribed *bona fide* off-target sites for all these editors in human cells. Most importantly, we leveraged ONE-seq to generate the first experimentally-derived population-scale off-target profiles for Cas9 nucleases that define the impacts of sequence variants from >2,500 individual human genome sequences in the 1000 Genomes Project database. Notably, some of the variants we identified that lead to increased mutation frequencies at off-target sites are enriched in specific human populations. We validated that novel population-specific, variant-sensitive off-target sites nominated by ONE-seq *in vitro* can show increased frequencies of mutations in human lymphoblastoid cells (**LCL**s) harboring these sequence variants. Collectively, our results demonstrate that ONE-seq is a highly sensitive off-target nomination method that can uniquely be used to identify population subgroup-linked differences in off-target profiles of gene editors. ONE-seq provides an important new pathway by which to assess the impacts of global human genetic sequence diversity on the specificities of gene editors, thereby enabling a broader and more all-inclusive approach for profiling off-target effects of these transformative therapeutic technologies.

## Background and Introduction

Gene-editing nucleases and CRISPR base editors are sequence-specific modification technologies that efficiently alter genomic targets of interest but that can also induce unwanted off-target mutations at sites resembling the on-target sequence. The most commonly used and widely accepted strategy for off-target determination of gene editors employs a two-step approach consisting of nomination and validation^1–10^. In the initial nomination step, a highly sensitive method is used to identify a superset of candidate off-target activity sites for a given editor. In a subsequent validation step, nominated sites are assessed in cells or organisms in which the gene editor is expressed to determine whether they show evidence of mutations. Therefore, sensitivity of the initial nomination step is of critical importance because it defines and limits the sites examined in the subsequent validation step. Various cell- or organism-based (e.g., GUIDE-seq^6^, BLESS^11^/BLISS^12^, HTGTS^13^, DISCOVER-seq^10^) and *in vitro* (e.g., Digenome-seq^5^, CIRCLE-seq^7^/CHANGE-seq^14^, SITE-seq^15^) nomination methods have been previously described. *In vitro* nomination methods generally exhibit higher sensitivity^7, 14, 16^ and have been used successfully to identify genomic sites that show *bona fide* off-target mutations in cells and organisms^1–5, 7^.

A significant limitation of all existing off-target nomination assays is that they can only be performed on one genome at a time, making it impractical to comprehensively assess the tremendous diversity of sequence variants that exist in large populations of humans or other organisms. Two *in silico* studies previously illustrated that sequence variants can fall within potential off-target sites of gene editors^17, 18^, but the degree to which these sequence differences might actually increase or decrease mutation frequencies at these sites was never experimentally demonstrated. Two additional studies illustrated anecdotally how individual sequence variants can influence mutation frequencies at CRISPR-Cas9 off-target loci: one assessed a single variant at a single site^19^ while the other used a modified version of CIRCLE-seq to assess the impacts of variants from seven individual genomes^14^. However, these studies only provide small surveys of single genomes and do not assess the impacts of the broad repertoire of sequence variants that exist in the global human population and that are being rapidly identified in greater numbers with increasingly larger-scale genome sequencing projects^20–22^. This is an important limitation to address given that therapeutics in development will ultimately be given to large numbers of genetically diverse patients or to those with diseases that are highly prevalent in specific populations (e.g., sickle cell disease, beta-thalassemia). Despite this critical need, to our knowledge no experimental approach currently exists that can readily and robustly assess the impacts of sequence variants at population scale during the course of identifying, optimizing, and ultimately administering these gene editing technologies.

## Results

### Overview of the ONE-seq assay

To create a robust, sensitive, and universal *in vitro* off-target nomination assay that could be used at population-scale with any gene-editing nuclease or base editor, we envisioned a strategy in which we would build libraries of potential off-target sites using high-throughput custom DNA synthesis instead of cell-derived genomic DNA (**Fig. 1a**, top panel). The use of synthetic DNA overcomes two important limitations of genomic DNA-based assays: First, genomic DNA libraries used in existing *in vitro* off-target assays (e.g., Digenome-seq, CIRCLE-seq, and SITE-seq; **Fig. 1a**, bottom panel) harbor a vast excess of unrelated DNA sequences that are not cleaved or modified by a given gene editor of interest (**Fig. 1b**); this leads to a higher background of random genomic sequence reads, which in turn reduces assay sensitivity and/or requires higher numbers of sequencing reads. Second, the use of genomic DNA makes existing assays difficult to perform with more than one genome at a time, limiting the abilities of these methods to assess the effects of SNPs from a large number of individuals at population scale.

**Figure 1.**
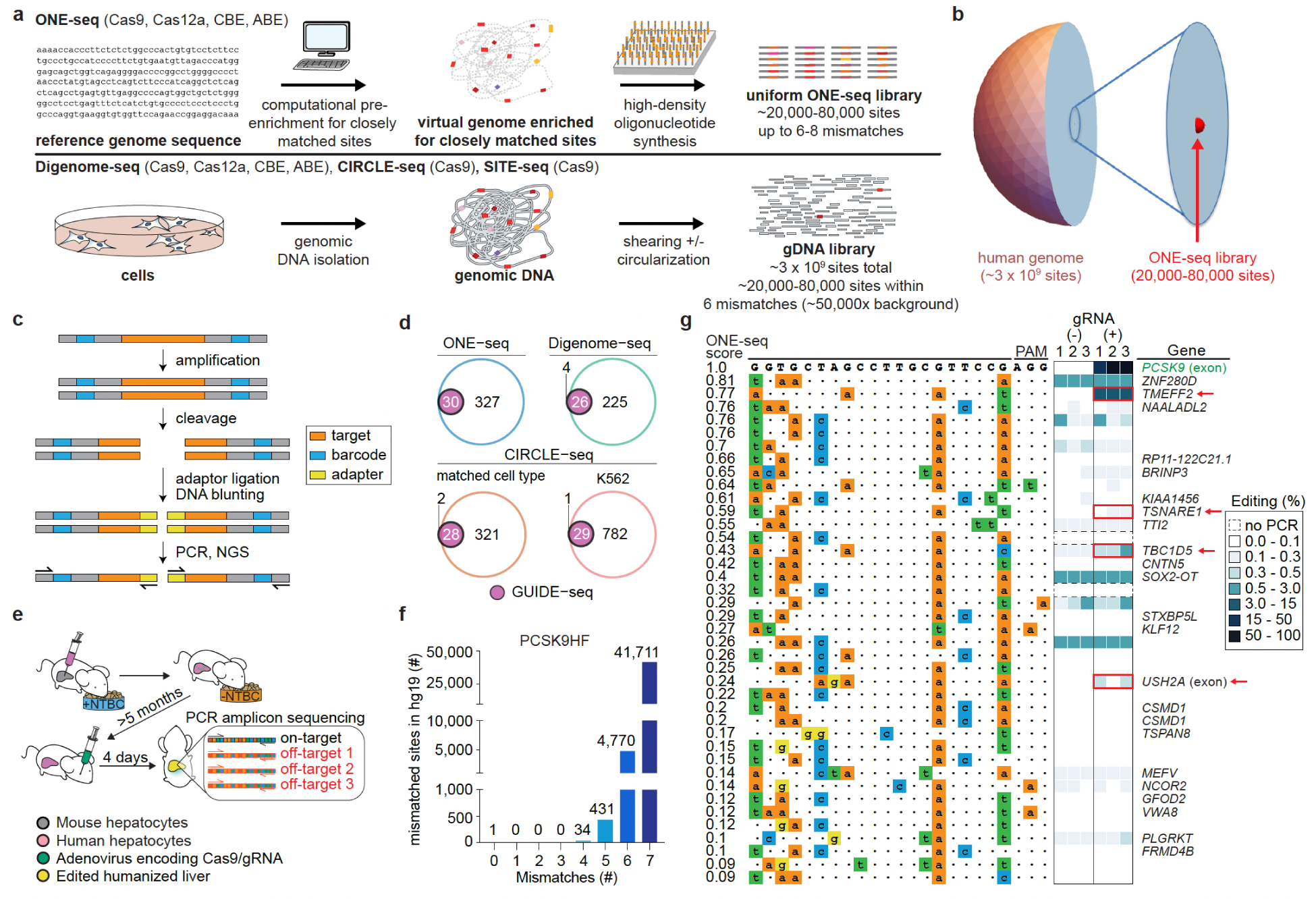
Overview of ONE-seq selections and profiling of Cas9 off-targets in human cells in culture and *in vivo*. **a,** Schematic comparing the workflow of ONE-seq (top) with existing *in vitro* methods (bottom). **b,** Comparison of number of sites in the human genome to those in a typical ONE-seq library. **c**, Schematic overview of ONE-seq selections with a gene-editing nuclease. **d**, Venn diagrams comparing abilities of ONE-seq, CIRCLE-seq, and Digenome-seq (open colored circles) to nominate *bona fide* off-target sites previously validated by GUIDE-seq (solid purple circles). **e**, Overview of the xenotransplant humanized liver mouse model system used to validate SpCas9 nuclease off-targets *in vivo*; NTBC denotes nitisinone. **f**, Graph showing the numbers of *in silico*-predicted sites in the hg19 reference genome with the indicated number of mismatches to the PCSK9HF target site. **g**, Testing of ONE-seq-nominated sites for the SpCas9 PCSK9HF gRNA from liver humanized mice using targeted amplicon sequencing. The heatmap shows indel frequencies and sites showing statistically significant frequencies of indels relative to a negative control are indicated by red boxes and red arrows.

In contrast to existing *in vitro* off-target assays that use genomic DNA, the OligoNucleotide Enrichment and sequencing (**ONE-seq**) method computationally enumerates all sites in one or more genomes of interest that harbor a specified number of differences (mismatches and/or DNA or RNA “bulges”) relative to the intended on-target site of a given gene editing nuclease or base editor (**Fig. 1a**, top panel; **Methods**). High-throughput oligonucleotide synthesis is then used to create a library of fixed-length ∼200 nt single stranded DNAs each harboring one of the computationally identified mismatched sites (**Fig. 1a** and **Extended Data Fig. 1a**).

Each site in the library is also embedded within a common DNA sequence context and associated with unique barcodes at both ends of the oligonucleotide that permit its unambiguous identification even after cleavage and/or modification (**Fig. 1c, Extended Data Fig. 1a**). Oligos are released from the solid support on which they are synthesized and then converted to double-stranded DNA by limited-cycle liquid-phase PCR (**Extended Data Fig. 1a**; **Methods**). The resulting pre-selection ONE-seq libraries can then be treated with a gene-editing nuclease or base editor of interest to identify off-target sites of activity as described in more detail below.

### Construction and characterization of ONE-seq libraries

To test the feasibility of the ONE-seq strategy, we first built libraries for 14 different *Streptococcus pyogenes* Cas9 (**SpCas9**) guide RNAs (**gRNA**s) and 4 different *Lachnospiraceae bacterium* Cas12a (**LbCas12a**) gRNAs (**Supplementary Table 1**). These 18 gRNAs target various sites in the human genome and were chosen because they have either been used in previous studies with base editors (ABE site 14, ABE site 16, ABE site 18; abbreviated here as ABE14, ABE16, ABE18)^23^ or had their gene-editor off-target profiles characterized previously (the other 15 gRNAs) using one or more existing methods^6–9, 24–26^.

For 13 of the 14 SpCas9 gRNAs, we synthesized ONE-seq libraries harboring all DNA sites in the hg19 reference human genome sequence that have up to six mismatches relative to the intended on-target site and all sites that have up to four mismatches and at least a one base DNA or RNA bulge (**Extended Data Figs. 1b and 1c**; **Supplementary Table 2**; **Methods**). For the VEGFA site 3 gRNA, which has an unusually large number of closely matched sites in hg19, we synthesized a library harboring sites with up to four mismatches and sites with up to two mismatches together with a single bp DNA or RNA bulge (library size of 23,955) (**Extended Data Fig. 1b**; **Supplementary Table 2**). For the four LbCas12a gRNAs, we synthesized libraries with sites from hg19 harboring up to eight mismatches relative to the intended on-target site, which we able to do because the longer LbCas12a target sites generally have fewer mismatched sites in the genome. The sizes of these 17 libraries (range 23,522 to 97,644; **Extended Data Fig. 1b**; **Supplementary Table 2**) were such that we could readily synthesize them using commercially available, high-density chip-based oligonucleotide synthesis (**Methods**). Characterization of these 18 libraries using next-generation sequencing (**Methods**) showed exceptionally high coverage of all synthesized sites (99.93% of all library members present among all 18 libraries; **Supplementary Table 3**) and high uniformity of sequence representation within each library with a mean 90/10 ratio of 2.5 +/-0.5 (**Extended Data Fig. 1d**; **Supplementary Table 3**). For 14 of the 18 ONE-seq libraries, the uniformity of mismatched site representation was superior to what was observed for these same sequences in a CIRCLE-seq library (**Supplementary Result 1**; **Extended Data Fig. 1d**; **Supplementary Table 3**).

### Validating ONE-seq for highly sensitive identification of bona fide Cas9 nuclease off-targets in human cells

We tested the efficacy of ONE-seq for identifying SpCas9 nuclease off-target effects in human cells by performing selections with five different gRNAs (*RNF2*, HEK293 site 2 [abbreviated as HEK2], HEK293 site 3 [abbreviated as HEK3], *FANCF*, and *EMX1*), each previously profiled for off-target activity by one or more existing off-target assays^6, 7, 24^. Following incubation of ONE-seq libraries with purified SpCas9 and the cognate gRNA, we captured resulting double-strand breaks by ligation of next-generation sequencing (**NGS**) adapters after an end-blunting step (**Fig. 1c**; **Methods**). Following gel extraction, amplification, and NGS, cleaved sites were counted based on the identities of flanking barcodes and each assigned a “ONE-seq nuclease score” normalized to activity at the on-target site (**Methods**). Results from these ONE-seq selections showed that the on-target site was the most highly enriched or among the most highly enriched library members, demonstrating the efficacy of the method for identifying cleaved sites (**Extended Data Fig. 2a).**

**Figure 2.**
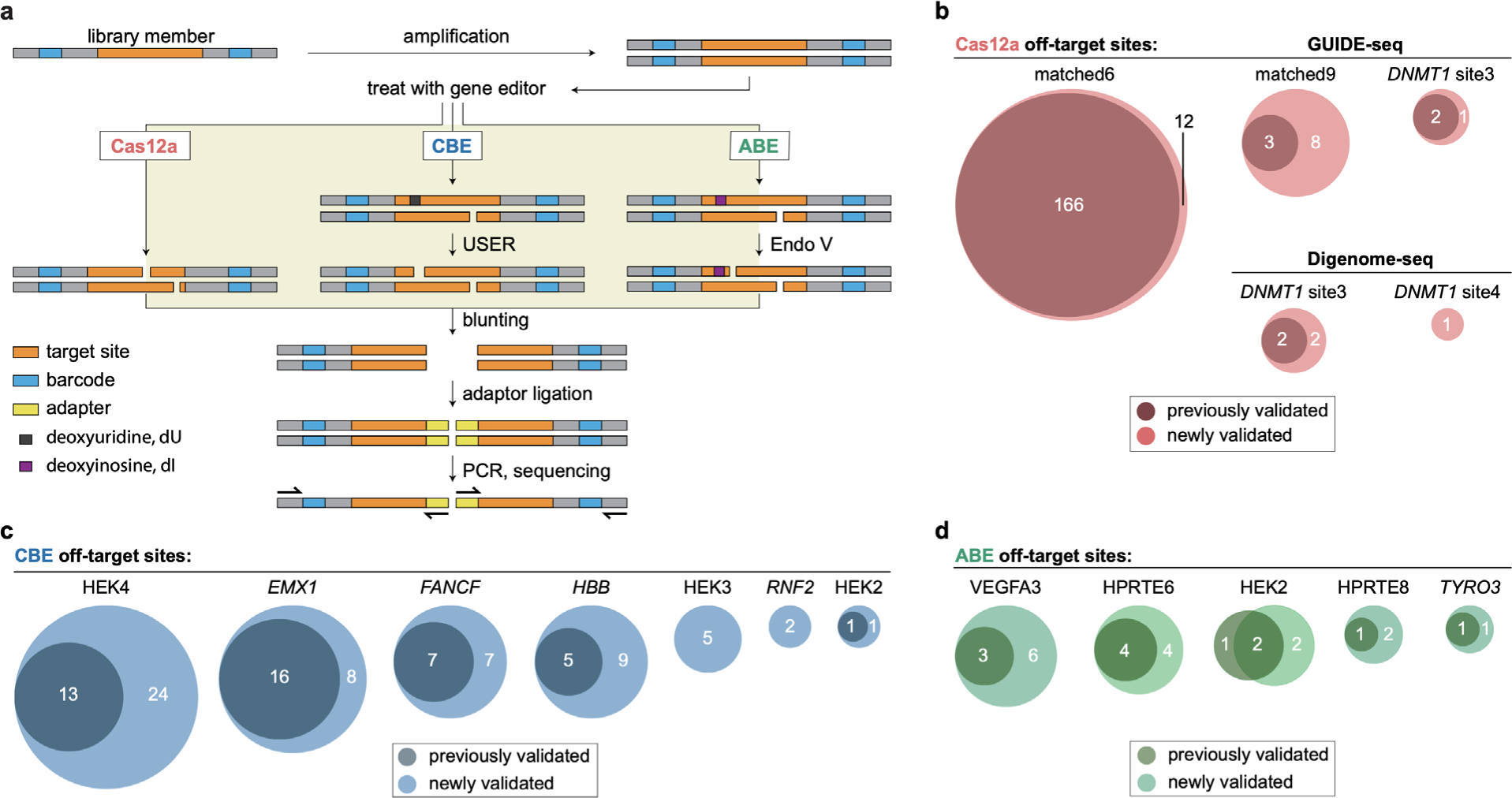
ONE-seq outperforms existing methods for nominating *bona fide* Cas12a, CBE, and ABE off-targets in human cells. **a,** Schematic overview of ONE-seq selections for Cas12a nucleases, CBEs, and ABEs. **b**, Venn diagrams illustrating identification of previously validated and newly validated LbCas12a off-target sites by ONE-seq; comparisons are shown for four different gRNAs previously assayed by GUIDE-seq or Digenome-seq. **c,** Venn diagrams illustrating identification of previously validated and newly validated CBE off-target sites by ONE-seq; comparisons are shown for seven different gRNAs previously assayed by Digenome-seq. **d**, Venn diagrams illustrating identification of previously validated and newly validated ABE off-target sites by ONE-seq; comparisons are shown for five different gRNAs previously assayed by Digenome-seq. **b-d,** All sites shown as validated by ONE-seq (light colored circles) had ONE-seq scores >0.01. CBE, cytidine base editor; ABE, adenine base editor.

To compare the relative sensitivity of ONE-seq with other existing *in vitro* nomination assays (Digenome-seq and CIRCLE-seq), we assessed their efficacies for nominating *bona fide* SpCas9 off-target sites (i.e., those that show evidence of indel mutations in human cells). Previously published experiments had identified 30 *bona fide* SpCas9 off-target sites for four of the five gRNAs described above in human U2OS cells or HEK293 cells (determined using the GUIDE-seq method)^6^. ONE-seq successfully nominated all 30 of these *bona fide* off-target sites, with all of them having ONE-seq scores > 0.01 and being among the most highly enriched ONE-seq candidates (**Fig. 1d; Extended Data Figs. 2a and 2b**). These results are comparable to those obtained with CIRCLE-seq, which also collectively nominated all 30 GUIDE-seq off-target sites across selections performed with genomic DNA from three different human cell lines (28 of the 30 *bona fide* off-target sites were found with CIRCLE-seq performed in matched U2OS or HEK293 cells and 29 of the 30 *bona fide* off-target sites in K562 cells) (**Fig. 1d** and **Extended Data Fig. 2b**). By contrast, Digenome-seq experiments nominated only 26 of the 30 *bona fide* off-target sites (**Fig. 1d** and **Extended Data Fig. 2b**). Importantly, ONE-seq also nominated 13 *bona fide* off-target sites additionally nominated by CIRCLE-seq (but not by GUIDE-seq) for the *EMX1* gRNA in an earlier study^7^ (**Extended Data Fig. 2a**). Taken together, these results show that ONE-seq is at least as sensitive as CIRCLE-seq and more sensitive than Digenome-seq and GUIDE-seq for nominating *bona fide* SpCas9 off-target sites in human cells.

### ONE-seq identifies bona fide Cas9 off-target sites in vivo

We also tested whether ONE-seq could identify *bona fide* off-target mutation sites *in vivo* in a clinically relevant animal model. To do this, we used humanized mice in which xenotransplanted human hepatocytes have replaced mouse hepatocytes in the liver^27^ (**Fig. 1e**; **Methods**), thereby enabling a more translationally relevant assessment of off-target mutations in liver cells with human (instead of mouse) genomes. To perform a highly rigorous test of ONE-seq, we used it to profile SpCas9 with a gRNA (hereafter referred to as the PCSK9HF (high-fidelity) gRNA) that targets a site in the human *PCSK9* coding sequence for which there are no closely matched sites in the hg19 reference human genome sequence with one to three mismatches relative to the intended on-target site (**Fig. 1f**). We performed ONE-seq selections with a library containing all sites from hg19 that had up to seven mismatches relative to the PCSK9HF on-target site and also all sites containing up to four mismatches and at least one DNA or RNA bulge. This selection nominated numerous off-target sites (**Extended Data Fig. 3**, **Supplementary Table 4**), and we then examined the on-target site and the 40 off-target sites with the highest ONE-seq scores in the livers of xenotransplanted human hepatocyte mice four days after they had been treated with adenovirus encoding for SpCas9 and PCSK9HF gRNA. These targeted amplicon sequencing experiments revealed efficient on-target editing of the *PCSK9* gene (mean indel frequency of 53.6%) and significant evidence of indel mutations at four of the 40 off-target sites (mean frequencies ranging from 0.1 to 6.8%) (**Fig. 1g**). Among the four mutated off-target sites, two had four mismatches and two had five mismatches relative to the on-target site (**Fig. 1g**). These results demonstrate the efficacy and high sensitivity of ONE-seq for identifying *bona fide* off-target sites in an *in vivo* setting for a Cas9 gRNA and nuclease pre-identified *in silico* to have no closely matched sites in the reference hg19 human genome.

**Figure 3.**
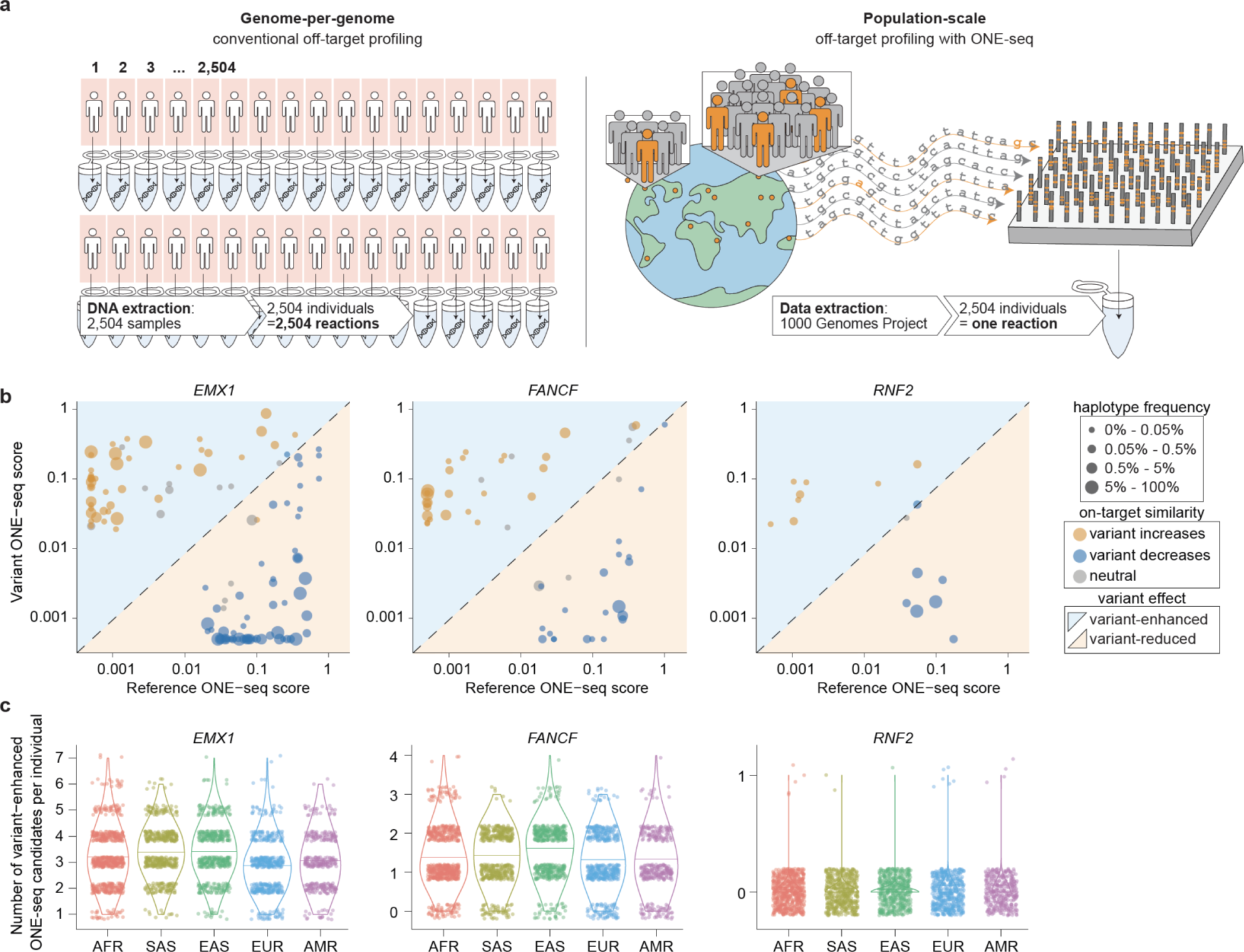
Population-scale, sequence variant-aware SpCas9 nuclease off-target profiling using ONE-seq. **a,** Schematic illustrating genome-per-genome off-target analysis used by previous methods (left panel) and population-scale, sequence-variant aware off-target profiling using ONE-seq (right panel). **b,** Scatterplots of matched variant and reference site ONE-seq scores for variants that lead to a statistically significant change of the ONE-seq score for three different SpCas9-gRNA nucleases. Orange and blue circles indicate off-targets for which genomic variants increase or decrease similarity to the intended on-target similarity, respectively. Circle sizes denote haplotype frequencies. Blue and orange background correspond to areas of the plot in which variant-enhanced or variant-reduced off-targets, respectively, would be present. **c,** Numbers of variant-enhanced off-target candidates per individual stratified by super-population. Each dot represents an individual from the 1000 Genomes Project. AFR, African; SAS, South Asian; EAS, East Asian; EUR, European; AMR, American.

### Identification of bona fide Cas12a nuclease off-targets with ONE-seq in human cells

Because many gene-editing nucleases leave overhangs rather than blunt ends (e.g., Cas12a/Cpf1, zinc finger nucleases, TALENs, homing endonucleases), we also assessed whether ONE-seq could be used to identify off-target cleavage sites for these other types of nucleases. We were particularly interested in doing so because our attempts to adapt CIRCLE-seq to include an extra end-blunting step had led to a greater than 10-fold reduction in usable sequencing reads, thereby rendering the assay ineffective and insensitive for nucleases that leave overhang ends (**Supplementary Result 2**). By contrast, ONE-seq includes an end-blunting step as part of its standard protocol (**Fig. 2a**; **Methods**). We tested ONE-seq with the LbCas12a nuclease and four different crRNAs (matched6, matched9, *DNMT1* site 3, and *DNMT1* site 4), which had been previously profiled for off-target sites using GUIDE-seq and/or Digenome-seq^25, 26^. These earlier studies had validated a total of 166, three, four, and no *bona fide* off-target sites for the matched 6, matched 9, *DNMT1* site 3, and *DNMT1* site 4 crRNAs, respectively, in human U2OS and/or HEK293T cells. ONE-seq successfully nominated all 173 of these *bona fide* off-target sites, with nearly all of them having amongst the highest ONE-seq scores (**Extended Data Figs. 4a)**.

**Figure 4.**
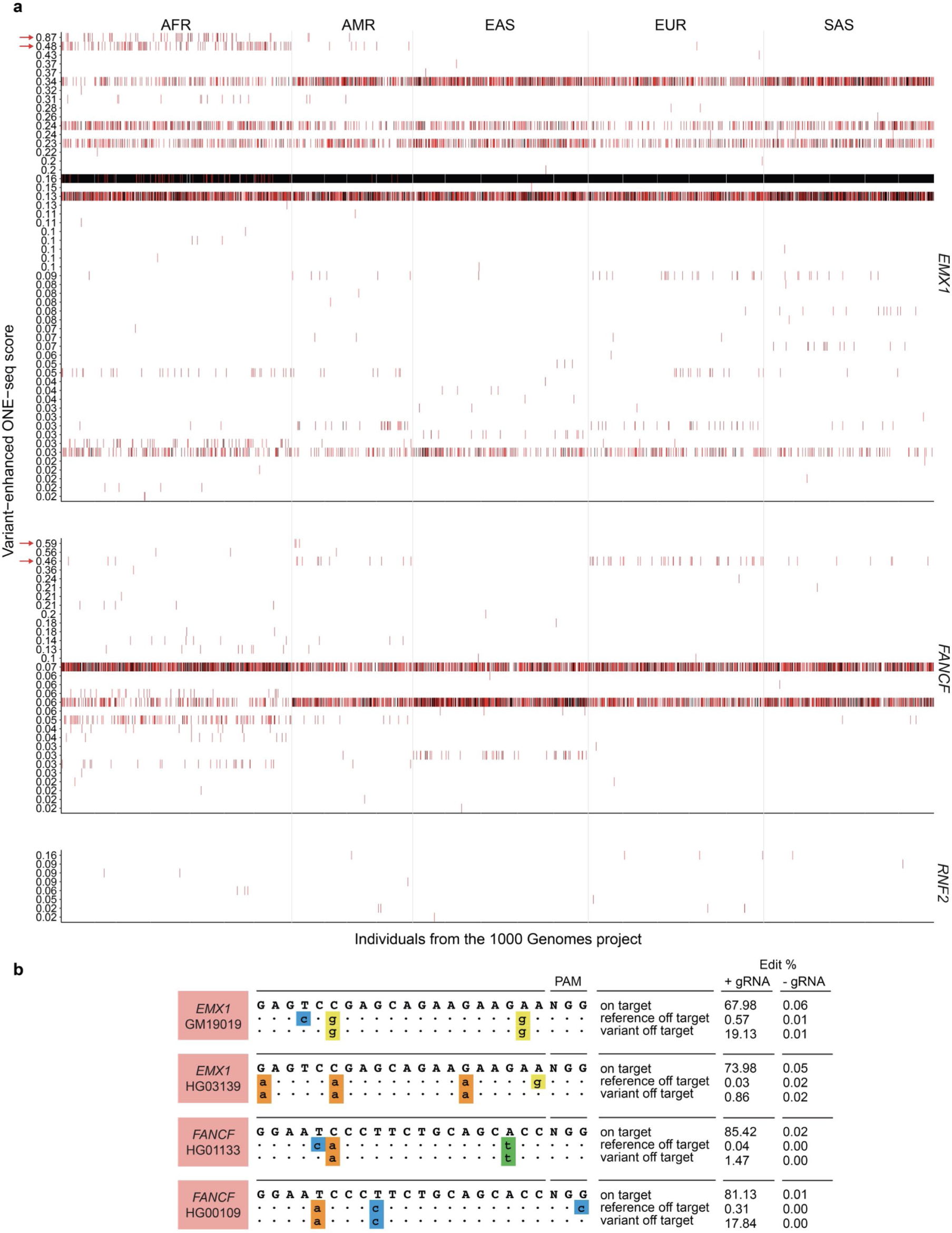
ONE-seq identifies population-specific sequence variants that enhance off-target cleavage *in vitro* and increase off-target mutation frequencies in human cells. **a,** Heatmap displaying statistically significant variant-enhanced off-target sites for three SpCas9-gRNA nucleases. Each column denotes an individual from the 1000 Genomes project, clustered by superpopulation as labeled at the top of each panel. Within each heatmap, variant sites are arranged by their ONE-seq nuclease scores (vertical axis labels). Red and black lines represent heterozygous and homozygous variant carrier status, respectively, for each individual, respectively. Variant sites selected for subsequent validation in human lymphoblastoid cell lines (LCLs) are marked with a red arrow. AFR, African; AMR, American; EAS, East Asian; EUR, European; SAS, South Asian. **b,** Results of targeted amplicon sequencing validation studies for selected variant-enhanced ONE-seq candidates in human LCLs. Indel frequencies for the on-target site, reference off-target site, and variant off-target site are shown (average of experimental triplicates). Lower case letters in colored boxes indicate mismatches relative to the on-target site. The target sgRNA and LCL identifier number are shown in the red box.

Because our Cas12a ONE-seq selections also nominated additional off-target cleavage sites that had not been found in previous Digenome-seq and GUIDE-seq experiments (**Extended Data Fig. 4a**), we performed two validations to examine whether these sites might also show evidence of indel mutations in human cells. In the first, to compare to GUIDE-seq^26^, which had been previously used with U2OS cells, we performed amplicon sequencing on genomic DNA of Cas12a nuclease-treated U2OS cells. We identified a total of 21 new *bona fide* off-target sites that had not been previously detected for the four gRNAs by GUIDE-seq: 12, 8, 1, and 0 new sites for the matched6, matched9, *DNMT1* site 3, and *DNMT1* site 4 gRNAs, respectively **(Extended Data Figs. 4a and 4b)**. Second, to compare to Digenome-seq^25^, which had been previously used with HEK293T cells, we performed amplicon sequencing on genomic DNA of Cas12a nuclease-treated HEK293T cells. We found a total of five *bona fide* off-target sites for *DNMT1* site 3 and *DNMT1* site 4 of which three sites had not been previously identified by earlier Digenome-seq studies (**Extended Data Figs. 4a and 4b**). Digenome-seq might have failed to detect some off-target sites because of limitations in its sensitivity^7^ (**Fig. 1d**; **Extended Data Fig. 2b)** and GUIDE-seq might have missed some off-target sites because integration of the GUIDE-seq dsODN tag in cells presumably requires filling in of overhangs at Cas12a nuclease-induced cut sites (which may not occur efficiently at all such sites). We also validated additional off-target sites in HEK293T cells that had been nominated using ONE-seq (**Extended Data Fig. 4b**). Taken together, our results demonstrate that ONE-seq outperforms Digenome-seq and GUIDE-seq with Cas12a nuclease, successfully identifying off-target sites that had not been identified by these other methods in earlier studies (**Fig. 2b**).

### Adaptation of ONE-seq to identify bona fide off-targets of CRISPR base editors in human cells

We additionally sought to apply ONE-seq to CRISPR cytosine base editors (**CBE**s)^28^ and adenine base editors (**ABE**s)^23^. To do this, we performed modified ONE-seq selections with the widely used BE3 CBE or the optimized ABE7.10, each with various gRNAs. In these reactions, DNA sites bearing a uracil (induced by deamination of cytosine by BE3) or an inosine (induced by deamination of adenine by ABE7.10) have a nick introduced at these base positions by treating with USER or Endonuclease V, respectively (**Fig. 2a**). The resulting nick together with another nick on the opposing strand induced by the Cas9 nickase activity of the base editor create a staggered end double-strand break (**DSB**), which can then be blunted and serve as a substrate for ligation of NGS adapters (**Fig. 2a**). Amplification, size selection, and sequencing of the resulting products enables identification of off-target sites by their associated unique flanking barcodes (**Fig. 2a**) and also calculation of a quantitative ONE-seq CBE or ONE-seq ABE score normalized to cleavage of the intended on-target site (**Methods**).

Using these modified ONE-seq selection methods, we assessed the BE3 CBE with eight different gRNAs (targeted to the *HBB*, HEK2, HEK3, HEK4, *RNF2*, *EMX1*, *FANCF*, and ABE18 sites) and ABE7.10 with nine different gRNAs (targeted to the ABE14, ABE16, ABE18, HEK2, HEK3, *VEGFA3*, *HPRTE6*, *HPRTE8* and *TYRO3* sites). We chose these gRNAs because other groups had previously used the Digenome-seq method to nominate off-target sites for seven of the eight gRNAs with BE3^8^ and five of the nine gRNAs with ABE7.10^9, 29^. Among the sites previously nominated by Digenome-seq, a total of 42 BE3 off-target sites among the seven gRNAs and a total of 12 ABE7.10 off-target sites among five of the nine gRNAs (HEK2, *VEGFA3*, *HPRTE6*, *HPRTE8*, *TYRO3*) were confirmed as *bona fide* sites of mutation in human HEK293T cells using targeted amplicon sequencing^8, 9, 29^. All 42 BE3 and 11 of 12 ABE7.10 off-target sites reported as validated in these previous studies had a ONE-seq score of >0.01 in our selections (**Extended Data Figs. 5a** and **6a**). Furthermore, all 54 of these off-target sites had among the highest ONE-seq CBE or ONE-seq ABE scores within their respective selections (**Extended Data Figs. 5a** and **6a**). Even the one ABE7.10 off-target site that had a ONE-seq score of <0.01 was enriched as the 103^rd^ ranked site out of the 39,474 mismatched sites present in the HEK2 gRNA library.

We next explored whether additional base editor off-target sites nominated by ONE-seq but not nominated or validated in previous Digenome-seq studies might also show evidence of editing in human cells. For each of the eight BE3 gRNAs and nine ABE7.10 gRNAs, we used targeted amplicon sequencing to assess ∼20-40 sites in HEK293T cells that were: (a) nominated by ONE-seq but not by Digenome-seq (**Type I sites**), (b) nominated by both ONE-seq and Digenome-seq but not shown to be edited in previous human cell-based validation experiments (**Type II sites**), or (c) nominated by ONE-seq for base editor/gRNAs that had not previously characterized by Digenome-seq (**Type III sites**) sites (**Extended Data Figs. 5a, 5c, 6a, **and** 6c**). Among the seven BE3 gRNAs previously assessed by Digenome-seq^8^, we validated 28 Type I and 28 Type II sites as *bona fide* off-target sites in HEK293T cells (mean mutation frequencies ranging from 0.03 to 21% and from 0.07 to 48%, respectively) (**Extended Data Figs. 5a and 5b**). For the eighth BE3 gRNA (ABE18) not previously assessed by Digenome-seq, we also validated six Type III sites as new *bona fide* off-target sites (mean mutation frequencies ranging from 0.24 to 9.6%) (**Extended Data Figs. 5c and 5d**). Among the five ABE7.10 gRNAs previously evaluated by Digenome-seq or a variant Digenome-seq method known as EndoV-seq^9, 29^, we validated one new *bona fide* Type I off-target site (for the VEGFA3 gRNA; mean mutation frequency of 0.6%) and nine new *bona fide* Type II off-target sites (mean mutation frequencies ranging from 0.06 to 0.6%) in HEK293T cells (**Extended Data Figs. 6a and 6b**). For the four other ABE7.10 gRNAs not previously assessed by Digenome-seq, we also validated five *bona fide* Type III off-target sites (mean mutation frequencies ranging from 0.05 to 19%) (**Extended Data Figs. 6c and 6d**). To further increase our sensitivity for identifying *bona fide* off-target sites, we also assessed the ONE-seq-nominated ABE sites for editing in HEK293T cells in which the ABE7.10 editor was expressed at much higher levels (identified by flow cytometry; **Methods**). These experiments generally yielded higher edit frequencies at on- and off-target sites for all sites examined but also validated seven additional *bona fide* off-target sites: two Type I sites, two Type II sites, and three Type III sites (**Extended Data Fig. 6**). Taken together, these data demonstrate that ONE-seq substantially outperforms Digenome-seq for nominating *bona fide* CBE and ABE off-target sites in human cells (**Figs. 2c and 2d**).

### Using ONE-seq to identify sequence variant-sensitive off-target sites at population scale

As noted above, in contrast to the “one-by-one“ nature of existing *in vitro* nomination methods that can only interrogate a single individual genome at a time (**Fig. 3a**, left panel), ONE-seq can assess the impacts of genetic variation from thousands of genomes simultaneously in a single reaction (**Fig. 3a**, right panel). This is possible with ONE-seq because each unique variant sequence only needs to be represented once in the *in vitro* selection library. To conduct an initial proof-of-concept, we created an informatics pipeline that analyzed genomes from 2,504 ethnically diverse individuals present in the 1,000 Genome Project database^20^ to identify all SNPs, insertions, or deletions that fall within all sites present in the original ONE-seq libraries we built for the SpCas9 *EMX1*, *FANCF*, and *RNF2* gRNAs (**Methods**). Using this output, we constructed new “variant-aware” ONE-seq libraries harboring both the 36,159, 20,043, and 15,686 sequence variant sites and matched sites from hg19 reference genome for the SpCas9 *EMX1*, *FANCF*, and *RNF2* gRNAs, respectively (**Supplementary Table 5**). Of note, the majority of such variants we found for all three gRNAs were unique to a single 1000 Genomes Project super population (**Extended Data Fig. 7**).

To identify “variant-sensitive” SpCas9 off-target sites (i.e., those for which the presence of a variant alters cleavage), we performed ONE-seq selections with the variant-aware libraries and then compared scores for matched reference/variant site pairs from these experiments. For all sites that were cleaved in each selection, we calculated ONE-seq nuclease scores normalized to the on-target site included in each experiment as a positive control (**Methods**). We found 121, 55, and 16 “variant-sensitive” sites from the *EMX1*, *FANCF*, and *RNF2* libraries, respectively, that had significantly different (p < 0.05 after multiple comparisons adjustment) ONE-seq scores relative to their matched cognate hg19 reference genome sites (**Methods**; **Fig. 3b**; **Supplementary Table 6**). These variant-sensitive sites include some that have increased or decreased sequence similarity to the intended on-target site and some that neither increase nor decrease sequence similarity to the on-target site (**Supplementary Table 6**). Notably, for all three selections, all but one of the 80 sites with increased similarity to the on-target were “variant-enhanced” (variant site ONE-seq score > reference site ONE-seq score) and all 90 sites with decreased similarity to the on-target site were “variant-reduced” (variant site ONE-seq score < reference site ONE-seq score) (**Fig. 3b**). Sites with neither increased nor decrease similarity to the on-target site did not predictably into one category or other – i.e., some were variant-enhanced while others were variant-decreased (**Fig. 3b**). These results confirm that our variant-aware ONE-seq selections can successfully identify variants whose presence significantly impacts cleavage frequencies at potential off-target sites within the reference human genome.

The data from our variant-aware ONE-seq selections provide the opportunity to examine the distribution of variant-enhanced off-target alleles among the genomes of 2,504 individuals of diverse genetic ancestry from the 1000 Genomes Project. We focused on variant-enhanced sites because of their greater concern for translational applications, and we undertook two different approaches to analyze the data: (1) We quantified the impact of genetic variation on the off-target profiles of individuals by quantifying the variant-enhanced off-target sites identified by our ONE-seq selections in the genomes of each of the 2,504 individuals. For example, for the EMX1 gRNA, the average individual had at least one newly nominated off-target site compared to the hg19 reference, while some individuals had as many as seven newly nominated off-target sites (**Fig. 3c**). Not surprisingly, more variant-sensitive sites were generally observed per individual for SpCas9 gRNAs that had higher numbers of off-target sites identified from ONE-seq selections performed with reference genome sequence (**Fig. 3c**). The same analysis performed at the more granular population level shows variants present in these smaller sets of individuals can also expand off-target space (**Extended Data Fig. 8**). (2) We assessed the impact of off-target genetic variation on the population level by quantifying the prevalence of each variant-enhanced site among all populations from the 1000 Genome Project. This analysis revealed that some sites are preferentially found in one or more super-populations or populations (**Fig. 4a**). In addition, some variants are found across all populations, while a small number appear to be “errors” in the hg19 reference sequence (variants present in a majority of individuals but not found in the reference) (**Fig. 4a**).

Finally, we sought to test whether variant-enhanced sites identified from our population-scale ONE-seq selections would also show evidence of increased mutagenesis in human cells. For these validation studies, we used human lymphoblastoid cell lines (**LCL**s) from the 1000 Genomes Project that harbored four variant-enhanced sites: two sites from the *EMX1* gRNA selection that were more prevalent in the African superpopulation; and two from the *FANCF* gRNA selection – one more prevalent in the Colombian population and another in both the American and European super-populations (**Fig. 4a**; red arrows). Each of the four LCLs we used is heterozygous for the variant-enhanced site with the other allele harboring the hg19 reference genome site, providing the opportunity to compare the frequencies of off-target mutations at both sites in the same cells. Following transfection of the LCLs with SpCas9 and the appropriate cognate gRNA (**Methods**), we performed targeted amplicon sequencing and found that all four variant-enhanced off-target sites showed increased indel mutation frequencies relative to their matched reference sites (**Fig. 4b)**. In two of the four cases, the presence of a sequence variant led to high-frequency off-target edits of >15%. Taken together, our results demonstrate that ONE-seq can assess the impacts of thousands of sequence variants on off-target profiles in one reaction tube and successfully identify at population-scale *bona fide* variant-sensitive off-target sites that are cleaved both *in vitro* and in human cells.

## Discussion

Our results demonstrate that ONE-seq provides a universal assay for nominating off-target sites of gene-editing nucleases and CRISPR base editors with unsurpassed sensitivity. For Cas9 nucleases, it exceeds or matches the sensitivities of existing *in vitro* (Digenome-seq and CIRCLE-seq) and cell-based (GUIDE-seq) off-target nomination methods and it can successfully identify *bona fide* off-target mutations *in vivo* in a human liver mouse model. For Cas12a nucleases, CBEs, and ABEs, ONE-seq outperforms all existing off-target nomination strategies, identifying *bona fide* mutations in human cells not found previously by these methods. In addition, the demonstration that ONE-seq works with Cas12a nucleases that leave overhangs at the cut site suggests that it should also work with other nucleases such as engineered homing endonucleases, zinc finger nucleases (ZFNs), and transcription activator-like effector nucleases (TALENs). Because ONE-seq can explicitly experimentally interrogate all genomic sites bearing up to 7-8 mismatches (for target sites with high genomic orthogonality) and including DNA or RNA bulges for sites with four or fewer mismatches, it also provides a superior alternative to all of the various existing computational nomination methods that rely only on *in silico* predictions and are known to have limitations in their predictive capabilities^30–34^. In addition, ONE-seq offers technical advantages of high reproducibility and scalability (critical properties for comparing results across different experiments and replicates).

The custom-build nature of ONE-seq provides flexibility to expand its capabilities as related technologies and our understanding of off-target effects continue to improve. Our work here has focused on off-targets in human genomes but the ONE-seq methods and software can be easily extended to any organism for which whole genome sequences are available. Using existing commercially available high-throughput methods, it should be straightforward to build ONE-seq libraries for essentially any Cas9 gRNAs. For example, we calculated ONE-seq library sizes for a set of 481 gRNAs targeting 24 therapeutically relevant genes^17^ and found that these ranged in size from 9,208 to 26,066 sites from the hg19 reference genome if one includes sites with up to 6 nucleotide and/or up to 2 bulge mismatches relative to the on-target site (**Extended Data Fig. 9**). With this level of library diversity, ONE-seq can already assess potential off-target site sequence space sizes that match those interrogated by CIRCLE-seq and GUIDE-seq, which impose the same degree of sequence-mismatch restrictions at the informatic analysis level. Although Digenome-seq does not use explicit informatic restrictions, its inability to nominate sites found by other *in vitro* methods (presumably due to the lack of sensitivity that results from its use of a whole genome sequencing as a screening method) substantially limits its utility. Furthermore, as the scale of custom oligonucleotide synthesis continues with its upward trajectory, this ever-improving capability can be easily incorporated to further increase the sequence space of mismatched sites that can be interrogated by ONE-seq.

The most important advance uniquely provided by ONE-seq is its capability to robustly assess how the rich diversity of sequence variation in the global human population can impact the off-target profiles of gene editors. Our experimental identification and validation of off-target loci that are enriched in specific human superpopulations and/or populations show that unwanted effects of gene editors are not always equal across all individuals; that is, individuals within certain populations may be more likely to possess certain SNPs that can increase off-target mutation frequencies. These findings highlight a substantial limitation of all existing off-target assays – their inability to readily assess off-targets for more than a single cell line, an individual reference genome, or a small number of genomes. Our findings show how ONE-seq could be used to account for genetic variation early in the therapeutics development process, ideally before substantial efforts are made to develop a particular candidate editor. Doing so will likely be important for all therapeutic gene editors but, in particular, when screening potential candidates for treating diseases that have a predominance in certain specific populations (e.g., sickle cell anemia, beta-thalassemia).

Although our proof-of-principle work here using 2,504 human genome sequences is substantially far more expansive and inclusive than any other previous off-target studies, many populations, including several with African genetic ancestry, still remain substantially underrepresented or missing from the 1000 Genomes Project data set. A more comprehensive accounting of the impacts of global human variation will require whole genome sequencing data for a much larger number of individuals. Fortunately, as noted above, the expandability of the ONE-seq approach makes it well positioned to readily take advantage of ever-expanding catalogues of human genome variation being made available in the public domain (e.g., the 1,000,000 genomes project, GnomAD). We believe that ONE-seq will provide a robust and tractable pathway to assess the potential impacts of global human genetic diversity on current and future gene editing therapeutic development efforts, thereby enabling a more expansive and inclusive approach for profiling the off-target effects of these transformative therapeutic technologies.

## Data Availability

High-throughput sequencing reads will be deposited in the NCBI Sequence Read Archive database after publication.

## Acknowledgements

This work was supported by the Defense Advanced Research Projects Agency (HR0011-17-2-0042 to M.J.A, L.P., and J.K.J). J.K.J. additionally received support from the U.S. National Institutes of Health (NIH) (RM1 HG009490 and R35 GM118158), the Desmond and Ann Heathwood MGH Research Scholar Award, and the Robert B. Colvin, M.D. Endowed Chair in Pathology. L.P. additionally received support from the U.S. NIH (R35 HG010717 and RM1 HG009490). K.M. is funded by the U.S. NIH (R35 HL145203) and the Winkelman Family Fund in Cardiovascular Innovation. D.R.L. is funded by U.S. NIH (U01 AI142756, RM1 HG009490, R35 GM118062) and the Howard Hughes Medical Institute. K.P. was funded by the Deutsche Forschungsgemeinschaft (DFG, German Research Foundation) – Projektnummer 417577129. G.A.N. was supported by the Helen Hay Whitney Fellowship. We thank Jason Gehrke, Julian Grünewald, Hana Kiros, and David Ma for helpful discussions and technical input and Ligi Paul-Pottenplackel for help with editing of the manuscript.

## Author Contributions

Wet lab experiments were performed by K.P., D.Y.K., K.E.S., H.S., H.M.L., J.A.G., and J.E.H.. K.P. and V.P. performed informatic analyses. K.P., M.J.A., J.K.J and V.P. wrote an initial draft of the manuscript and all authors contributed to the writing of the final version of the manuscript. G.A.N. and D.R.L. gave technical and conceptual advice and provided purified BE3 and ABE protein. K.C. and L.P. analyzed WGS data and nuclease amplicon sequencing. X.W. and K.M. performed experiments in xenotransplanted mice. M.C.C. analyzed 1000 Genomes Project data and designed variant-aware ONE-seq libraries. S.P.G analyzed CIRCLE-seq data. S.I. estimated ONE-seq library sizes. M.J.A. advised on statistical analyses. J.K.J. and V.P. supervised all research efforts on this project.

## Methods

### General methods

All chemicals were purchased from Sigma Aldrich and all enzymes were purchased from NEB unless otherwise stated. All DNA amplifications were conducted by PCR with Phusion High Fidelity DNA Polymerase (NEB) unless otherwise stated. Sequences of oligonucleotides are listed in the supplementary information (**Supplementary Table 7**). Sequences of transfection and protein expression plasmids used in this study can be found in the supplementary information (**Supplementary Table 8**).

### Cloning of gRNA expression plasmids and *in vitro* transcription of gRNA

gRNA expression plasmids for transfections were constructed by ligating annealed oligonucleotide duplexes into MLM3636 (without mCherry) or KESsgRNAmCherryBackbone(with mCherry) for Cas9/CBE/ABE experiments and BPK3082 for Cas12a experiments (U6 promoter human gRNA expression vectors). gRNA plasmids for *in vitro* transcription (IVT plasmids) were constructed by ligating annealed oligonucleotide duplexes into MSP3485 for Cas9/CBE/ABE experiments and RTW2675 for Cas12a experiments (T7 promoter bacterial gRNA expression vectors). For Cas9/CBE/ABE experiments, if a gRNA did not already have a 5’ guanine nucleotide (ex. HBB and ABE18), the 5’ nucleotide of the gRNA was substituted with a guanine. gRNAs were transcribed *in vitro* from the HindIII-digested IVT plasmids using the T7 RiboMAX Express Large Scale RNA Production System (Promega) according to manufacturer’s protocol with overnight incubation. gRNA was purified with the MEGAclear Transcription Clean-Up Kit (Thermo Fisher) according to manufacturer’s protocol.

### Generation of ONE-seq libraries

We used Cas-Designer^35^ (downloaded 2017-07-19) to search the human reference genome (hg19) for closely-matched sites for a given gene editor of interest. Depending on analyzed gRNA and gene editor, up to 6-8 mismatches to the on-target site were included in pre-selection libraries (**Supplementary Table 2**). Sequences with up to 2 DNA bulges and 4-6 mismatches and up to 2 RNA bulges and 3-6 mismatches were also included in the pre-selection libraries (**Supplementary Table 2**). The sequences of the closely-matched site plus additional 10 base pairs on each side of the closely-matched site were extracted using bedtools v2.25.0^36^. A unique 14 base pair barcode was assigned to each library member to facilitate post-selection identification. Barcodes were generated using a custom python script, and each barcode was required to have a Hamming distance of 2 or greater from every other barcode to reduce the risk of erroneous barcode readout caused by DNA sequencing or synthesis errors. The closely-matched sites including flanking DNA on each side were combined with barcode sequences with intervening constant sequences, constant3 and constant4, in a defined pattern (**Supplementary Table 9**). Additional constant sequences, constant2 and constant5, were attached to flank the barcodes and to serve as primer binding sites for library amplification. ONE-seq barcodes were uniquely assigned within the project to decrease the risk of cross contamination between different ONE-seq libraries. DNA sequences of ONE-seq libraries were synthesized on high-density oligonucleotide chips (Agilent Technologies; G7238A, G7222A). Oligonucleotide libraries were made double stranded by limited cycle PCR-amplification with primers oKP535 and oKP536/oKP577 (**Supplementary Table 7**), which contain flaps that additionally incorporate 8 or 11 base pair unique molecular identifiers, UMI1 and UMI2, and additional constant sequences, constant1 and constant6, (**Supplementary Table 9**).

For the creation of variant-aware ONE-seq libraries, reference-based ONE-seq libraries were created for EMX1, FANCF and RNF2 using cas-designer^35^ with up to 6 mismatches or up to 2 DNA bulges and 4 mismatches and up to 2 RNA bulges and 3 mismatches. The 1000 Genomes Project dataset was searched for positions non-structural genomic variants — including SNPs, insertions and deletions— that intersected with the set of positions from closely-matched sites identified by cas-designer. Since closely-matched sites with up to six mismatches to the on-target site were included in the reference-based ONE-seq library, variant sites with up to seven mismatches were included in the ONE-seq library if a genomic variant introduced an additional mismatch into a six mismatch site. Conversely, genomic variants that increased the on-target similarity could reduce the mismatch count of a six mismatch site to five mismatches. However, genomic variants transforming a seven mismatch site into a six mismatch site were not included in the libraries since the seven mismatch site was not part of the original reference-base ONE-seq library. Variants resulting in PAM creation (e.g. NGT > NGG) and PAM destruction events (e.g. NGG > NGT) were also detected by this approach, if they met the overall site mismatch parameters described above. Individual, population and super population haplotype frequencies were annotated for every variant from the haplotype-phased VCF files from the 1000 Genomes Project (phase 3) release. Phasing of off-target candidates that contained more than one variant were also performed in this step. Then, for every genomic variant or variant combination intersecting with a reference closely-matched site for a gRNA, the corresponding variant closely-matched site was constructed informatically by replacing, deleting or inserting nucleotides within the reference closely-matched sequence at the genomic variant position for SNPs, deletions or insertions, respectively. 10 bp of flanking genomic DNA were extracted using the local sequence of the variant containing haplotypes and the 43 bp variant sequence was embedded into the canonical ONE-seq library architecture including assignment of a ONE-seq library barcode (**Supplementary Table 9**). To set up a framework in which the influence of genomic variants or variant combinations on *in vitro* editing activity can be comparatively evaluated, the reference closely-matched site was always synthesized on the same chip as the inferred variant closely-matched site to allow the simultaneous assessment of variant/reference pairs of closely-matched sites. To allow ONE-seq score calculation the on-target site was included in each of the three variant-aware ONE-seq libraries.

### BE3 and ABE Protein Purification

Rosetta2 (DE3)-competent *E. coli* cells (MilliporeSigma) were transformed with plasmids encoding BE3 (pHR041^37^), or ABE7.10 (pGAN300^23^). Both editors were fused to a His6 N-terminal purification tag. Transformed colonies were grown overnight in 2xYT media containing 50 μg/ml kanamycin at 37 °C. The cells were diluted 1:125 into 1 liter of the same media and grown at 37 °C until OD_600_=0.70–0.8, about 3 hours. Cultures were cold-shocked on ice for 1 hour and protein expression was induced with 1 mM IPTG (GoldBio). Culture induction was continued for 16 hours 18 °C while shaking. Cells were collected by centrifugation at 6,000*g* for 20 min and resuspended in a final volume of 40mL cell lysis buffer (100 mM Tris-HCl, pH 8.0, 1 M NaCl, 20% glycerol, 5 mM TCEP (GoldBio), and 2 cOmplete™ EDTA-free protease inhibitor cocktail tablets (Sigma)). Cells were lysed by sonication on ice (10 min total using 3 s on 3 s off cycles) and the lysate cleared by centrifugation for 30 minutes at 20,000*g*. The cleared lysate was incubated with 1mL His-Pur nickel-NTA resin (Thermo Fisher) with rotation at 4 °C for 1 hour. The resin was collected by gravity-flow through an Econo-Pac chromatography column (Bio-Rad), then washed with 24 mL of cold lysis buffer. Bound protein was eluted with 2 mL cold elution buffer (100 mM Tris-HCl, pH 8.0, 0.5 M NaCl, 20% glycerol, 5 mM TCEP, 200 mM imidazole). The resulting protein fraction was further purified on a 5 ml Hi-Trap HP SP (GE Healthcare) cation exchange column using an Akta Pure FPLC. The nickel-NTA elution was diluted 25-fold in low salt buffer (100mM Tris-HCL, pH 8.0, 20% glycerol, 5mM TCEP) directly before loading on the column. After loading, the column was washed in 15mL low-salt buffer. The NaCl concentration was then increased in a linear gradient over 50 mL from 0 to 1M NaCl and collected in 1mL fractions. Protein-containing fractions were concentrated using an Amicon Ultra centrifugal filter with a 100,000 kDa cutoff (MilliporeSigma) centrifuged at 3,000*g.* Protein concentration was quantified using a reducing agent-compatible BCA assay (Pierce Biotechnology), following which aliquots were snap-frozen in liquid nitrogen and stored at -80°C.

### ONE-seq

ONE-seq libraries from Agilent Technologies were resuspended in 10 mM tris buffer and amplified in a 100 µL reaction containing 2 µL of 5 nM library, 1X Thermopol Buffer (NEB), 2.5 units of Taq polymerase, 200 µM dNTP, 0.2 µM of primers KP_extension_new_fw and KP_extension_new_rev or oKP577 (Supplementary Table 9). PCR reactions were purified with paramagnetic beads. For mixed library assays, two amplified ONE-seq libraries were mixed in a 1:1 ratio, and 30 ng of the mixed ONE-seq library was inputted into the *in vitro* cleavage or deamination reaction. *In vitro* deamination assays were performed in 100 µL reactions with CutSmart Buffer (1X), 30 ng of ONE-seq library, and an RNA:Base Editor:DNA ratio of 20:10:1 for BE3 and 200:100:1 for ABE. *In vitro* SpCas9 and Cas12 cleavage reactions were performed in 100 µL reactions with 1X Cas9 Reaction Buffer or NEBuffer 3.1 (NEB) for Cas9 or 1x NEBuffer 2.1 (NEB) for Cas12, 30 ng of library, and a RNA:Cas9:DNA ratio or RNA:Cas12:DNA ratio of 20:10:1. The RNA, buffer, and enzyme were mixed and pre-incubated at 25°C for 10 min before the amplified oligonucleotide library was added. The BE3, Cas9, and Cas12a reactions were incubated for 2 hours at 37 °C. The ABE reactions were incubated for 8 hours at 37 °C. After incubation, all the reactions were treated with 10 µL of Proteinase K (800 U/ml) at 37 °C for 10 min and the DNA was purified with paramagnetic beads. After purification, BE3 reactions were treated with USER enzyme (NEB) for 1 hour at 37°C in a 50 µL reaction, and ABE reactions were treated with Endonuclease V (NEB) for 30 min at 37 °C in a 20 µL reaction. Products of USER and Endonuclease V reactions were purified with paramagnetic beads. Cleaved products for BE3, Cas9, and Cas12a selections were extended at 72 °C for 10 min in a 50 µL reaction containing 1X Phusion HF Buffer, 200 µM dNTPs, 5% dimethyl sulfoxide (DMSO), and 1 unit of Phusion DNA polymerase (NEB). Cleaved products for ABE selections were extended at 37 °C for 30 min in a 50 µL reaction containing 1X NEBuffer 2, 200 µM dNTPs, and 15 units of Klenow Fragment (3’→5’ exo-, NEB). Following extension reactions, the products were purified with paramagnetic beads. For ABE selections, samples were additionally repaired, 5′ phosphorylated, 3′ dA-tailed using NEBNext® Ultra™ II End Repair/dA-Tailing Module (NEB E7546L) and were purified with paramagnetic beads. All selection samples were ligated to DNA adapters using the Quick Ligation kit (NEB) following the manufacturer’s protocol. Adapters were produced by annealing oKP145B and oKP146B or IVS_phosadaptABETOP and IVS_adapt_ABE_bottomT (Supplementary Table 9) in 1X STE Buffer (10 mM Tris-HCl pH 8.0, 1 mM EDTA, 100 mM NaCl). Adapter-ligated samples were purified by agarose gel electrophoresis and extraction with the QIAquick Gel Extraction kit (Qiagen) according to manufacturer’s protocol. For Cas9, Cas12a and BE3 selections, gel-purified products were amplified in 50 µL PCR reactions containing 12 µL of sample, 1X Phusion HF Buffer, 200 µM dNTPs, 1 unit of Phusion DNA Polymerase (NEB), 0.5 µM KP_P1 and 0.5 µM oKP101 for the PCR priming off region constant1 (**Supplementary Table 9**), henceforth called post-edit PCR1, or 0.5 µM oKP154 (Cas9/BE3)/oKP601(Cas12) for the PCR priming off region constant6 (**Supplementary Table 9**), henceforth called post-edit PCR2. For ABE selections, gel-purified products were amplified in 50 µL amplification reactions containing 12 µL of sample, 1X Thermopol Buffer, 200 µM dNTPs, 0.2 µM KP_P1, 0.2 µM oKP101, and 1.25 units of Taq DNA Polymerase (NEB). The products of the PCR reactions were purified with paramagnetic beads. A barcoding PCR was performed in 50 µL reactions containing 10 µL of selection product, 1X Phusion HF Buffer, 1 unit of Phusion DNA polymerase (NEB), 200 µM dNTPs, and 1 µM of each unique forward and reverse Illumina barcoding primer pair. After purification with paramagnetic beads, the samples were subjected to MiSeq or NextSeq sequencing.

### Mammalian cell culture and flow sorting

HEK293T cells were cultured in DMEM supplemented with 10% heat-inactivated fetal bovine serum, 2mM GlutaMax (ThermoFisher), penicillin, and streptomycin at 37°C and 5% CO_2_. U2OS cells were cultured in DMEM supplemented with 10% heat-inactivated fetal bovine serum, penicillin, and streptomycin at 37°C and 5% CO_2_. All lymphoblastoid cell lines (LCL) were cultured in RPMI supplemented with 15% heat-inactivated fetal bovine serum, 2mM GlutaMax (ThermoFisher), penicillin, and streptomycin at 37 °C and 5% CO_2_. Cell line identity was validated by STR profiling (ATCC) and the cultures were tested regularly for mycoplasma contamination. For ABE experiments without cell-sorting and BE3 experiments, 250,000 HEK293T cells were seeded per well in a 6-well plate and transfected 18 hours later. For ABE, 1.825 µg of pABE7.10 (Addgene Accession ID: #102919) and 675 ng of plasmid expressing gRNA were transfected using 7.5 µL of TransIT-X2 (Mirus) according to manufacturer’s protocol. For BE3, 1.825 µg of pJUL576 and 675 ng of plasmid expressing gRNA were transfected using the same protocol as for ABE. Genomic DNA was extracted 72 hours post-transfection with the QIAamp DNA Mini Kit (Qiagen) according to manufacturer’s protocol. For ABE experiments with cell-sorting, 3,100,000 HEK293T cells were seeded in a 10cm dish and transfected 18 hours later. For ABE, 7.5 µg of pJUL1459 and 2.5 µg of plasmid expressing gRNA were transfected using 45 µl of TransIT-X2 according to manufactureŕs protocol. 72 hours post-transfection, cells were sorted by flow cytometry selecting the ∼25% of cells with the highest GFP expression. After sorting, DNA was immediately extracted using the QIAamp DNA Mini Kit (Qiagen) according to manufacturer’s protocol. For Cas12a experiments with HEK293T, 3,100,000 HEK293T cells were seeded in a 10cm dish and transfected 18 hours later. 10 µg of SQT1665 (Addgene Accession ID: #78744) and 5 µg of plasmid expressing gRNA were transfected using 45 µl of TransIT-X2 according to manufactureŕs protocol. Genomic DNA was extracted 72 hours post-transfection with the QIAamp DNA Mini Kit (Qiagen) according to manufacturer’s protocol. For Cas12a experiments with U2OS, 4,000,000 U2OS cells were seeded in a 15cm dish and nucleofected 18 hours later. 1,000,000 cells were nucleofected with 3.33 µg of SQT1665 and 1.66 µg of plasmid expressing gRNA using SE Cell Line 4D-Nucleofector^TM^ X Kit L (Lonza) according to manufacturer’s protocol. Genomic DNA was extracted 72 hours post-transfection with the QIAamp DNA Mini Kit (Qiagen) according to manufacturer’s protocol. For all Cas9 experiments using lymphoblastoid cells, 20,000,000 lymphoblastoid cells were seeded in multiple T-25 flasks at a density of 400,000 cells/ml and were nucleofected 18 hours later. For treated samples, 10,000,000 lymphoblastoid cells were nucleofected with 60 µg of RTW3027 and 20 µg of plasmid expressing gRNA and mCherry using Cell Line NucleofectorTM Kit V (Lonza) according to manufacturer’s protocol. For control samples, 10,000,000 lymphoblastoid cells were nucleofected with 60 µg of RTW3027 using the same protocol as for treated samples. After 72 hours, cells were sorted by flow cytometry, selecting for all treated double positive Cas9-GFP/gRNA-mCherry expressing cells and all control Cas9-GFP expressing cells. Following cell sorting, DNA was immediately extracted using the QIAamp DNA Mini Kit (Qiagen) according to the manufacturer’s protocol.

### Deep sequencing of PCR amplicons

PCR primers were designed to yield a ∼70-270 bp product with the on-target or potential off-target site in the middle of the amplicon. On-target and potential off-target sites for BE3 and ABE, detected by ONE-seq, were amplified with PhusionU Multiplex PCR Master Mix (Thermo) in a 50 µL reaction with 100 ng input DNA. For Cas9 and Cas12a, on-target and potential off-target sites detected by ONE-seq were amplified with Phusion High Fidelity DNA Polymerase in a 50 µL reaction with 100 ng input DNA. On-target and SNP-induced off-target sites, detected by ONE-seq, were amplified with Phusion High Fidelity DNA Polymerase in multiple 50 µL reactions with 200 ng input DNA for on-target sites and 800 ng total input DNA for off-target sites. The PCR products were purified with paramagnetic beads. Product purity was assessed via capillary electrophoresis on a QIAxcel instrument (Qiagen). The NEBNext Ultra II DNA Library Prep kit (NEB) was used according to manufacturer’s protocol to ligate TruSeq (CD, formerly TruSeq HT) dual index adapters (Illumina Adapter Sequences, Document # 1000000002694 v09) to the PCR amplicons. The products were purified with paramagnetic beads and pooled according to concentrations measured using QuantiFluor® dsDNA System or droplet digital PCR (ddPCR). The pooled, adapter ligated, and indexed library was quantified again with ddPCR and sequenced with 2x150 bp paired end reads on an Illumina MiSeq sequencer.

### Xenotransplant model of human hepatocytes in mouse

All procedures used in animal studies were approved by the pertinent Institutional Animal Care and Use Committees at Harvard University, University of Pennsylvania, and Yecuris Corporation and were consistent with local, state, and federal regulations as applicable. The procedures were as previously described^27^. *Fah^−/−^Rag2^−/−^Il2rg^−/−^* (FRG KO) breeder mice on the C57BL/6 background were obtained from Yecuris Corporation. Mice were maintained on NTBC (also called nitisinone; Yecuris) prior to transplantation according to the manufacturer’s instructions. Twenty-four hours prior to transplantation, mice that were one to three months of age underwent intraperitoneal injection with 1 × 10^9^ pfu of adenovirus expressing the secreted form of urokinase-type plasminogen activator (Yecuris). For transplantation, 1 × 10^6^ primary hepatocytes (HEP10 Pooled Human Cryopreserved Hepatocytes; Thermo Fisher Scientific) were injected into the lower pole of the spleen.

During the surgery, 1%-2% inhaled isoflurane was used for anesthesia, and 0.05-0.1 mg/kg subcutaneous buprenorphrine was used as needed for analgesia in the perioperative and postoperative periods. Following transplantation, NTBC was gradually withdrawn over several weeks according to the manufacturer’s instructions, and human albumin levels in the blood were monitored on a monthly basis using the Human Albumin ELISA Quantitation Set (Bethyl Laboratories) according to the manufacturer’s instructions. Chimeric liver-humanized mice that were 8 to 11 months of age (at least 5 months following transplantation) were used for experiments. Mice were administered 1 × 10^11^ particles each via retro-orbital injection. 1%-2% inhaled isoflurane was used for anesthesia at the time of the injections. Three mice were given CRISPR-PCSK9 virus, and three mice were given CRISPR-control virus. As much as possible, the mice in the two groups were matched with respect to age and human albumin levels. After four days, the mice were euthanized by carbon dioxide asphyxiation. Whole liver samples were harvested for DNA analysis.

### CIRCLE-seq

CIRCLE-seq assays were conducted as previously described^7^ unless otherwise stated. For blunting experiments, Cas9-treated CIRCLE-seq libraries were blunted in a 50 µL reaction with 1X Phusion HF Buffer, 200 µM dNTPs, and 1 unit of Phusion HF polymerase (NEB) at 72 °C for 5 minutes. Following blunting, CIRCLE-seq protocol was continued with A-tailing and adaptor ligation. CIRCLE-seq data was analysed using hg19(GRCh27) and v1.1b of the CIRCLE-seq pipeline^7^ with the following parameters: window_size: 3, mapq_threshold: 50, start_threshold: 1, gap_threshold: 3, mismatch_threshold: 7, merged_analysis: False, variant_analysis: True.

### Data analysis

Amplicon sequencing data was analyzed with custom scripts employing R, Python and CRISPResso2^38^. Nuclease amplicon sequencing data was analysed with CRISPResso2 using default parameters for Cas9 and with ‘--quantification_window_center -5’ and ’--quantification_window_size 3’ to adapt the editing window for Cas12. For analyses of heterozygous lymphoblastoid cell lines both variant and reference allele were supplied as comma-separated string to CRISPResso2 as the ‘--amplicon_seq’ parameter. For nuclease experiments, reads were defined as edited if they contained an insertion or deletion in the specified editing window (CRISPResso default editing window for Cas9). Statistical significance was evaluated using Fisher’s exact test (for tests using python the ‘fisher_exact’ function of the ‘scipy.stats’ module was employed) and correction for multiple comparison was performed using the Benjamini-Hochberg method (for tests using python the ‘multipletests’ function of the ‘statsmodels.stats.multitest’ module was used with the argument “method=’fdr_bh’ ”). Additionally, to declare a tested genomic locus as edited, the average editing frequency of the gene editor treated samples were required to be at least three-fold higher than the average editing frequency of the control samples.

ONE-seq sequencing data were analyzed by custom Python scripts. Algorithms used are summarized in **Supplementary Algorithm**. Briefly, from the sequencing data, ONE-seq library member barcodes were read out to identify edited ONE-seq library members. Barcodes were counted and a ONE-seq score was calculated for each ONE-seq library member serving as a measure of *in vitro* cleavage or editing activity. The ONE-seq score for a library member was calculated as the ratio of sequencing read counts of the library member and the sequencing read counts of the ONE-seq library member representing the genomic on-target site for a given gene editor.

Variant-aware ONE-seq experiments were performed in triplicate and significance of within-replicate ONE-seq score differences between variant and reference sites were evaluated using a paired t-test (test performed in python using the ‘ttest_rel’ function of the ‘scipy.stats’ module) and Benjamini-Hochberg correction for multiple comparison (test performed in python using the ‘multipletests’ function of the ‘statsmodels.stats.multitest’ module with the argument “method=’fdr_bh’ ”). On-target alignment scores were calculated using the pairwise2 module of Biopython version 1.74. Upset plots were created using UpSetR^39^.

---

**Extended Data Fig 1.**
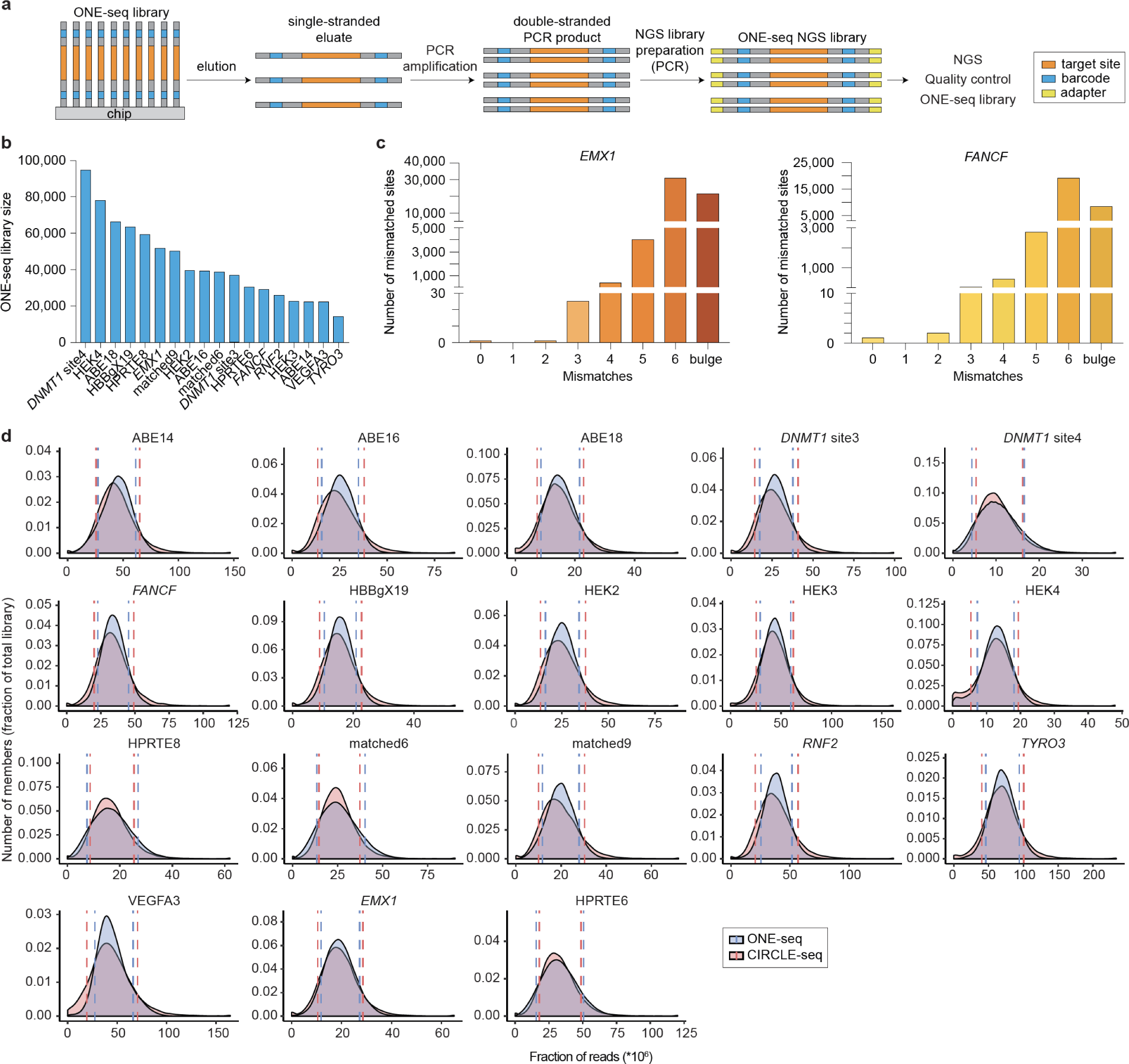
ONE-seq library generation and quality control. **a,** Schematic illustrating additional details of large-scale oligonucleotide chip synthesis, ONE-seq library amplification and ONE-seq library quality control via NGS. **b,** Graph showing ONE-seq library sizes for 18 SpCas9 or LbCas12a gRNAs used in this study. **c,** Graph showing distribution of library members for two representative ONE-seq oligonucleotide libraries (for the *EMX1* and *FANCF* SpCas9 gRNAs). The ONE-seq libraries shown contain genomic sites with up to 6 mismatches to the on-target site, including sites with DNA or RNA bulges. **d,** Density plots showing the coverage of ONE-seq library members in either ONE-seq (blue) and CIRCLE-seq (circle) libraries. NGS, next generation sequencing

**Extended Data Fig 2.**
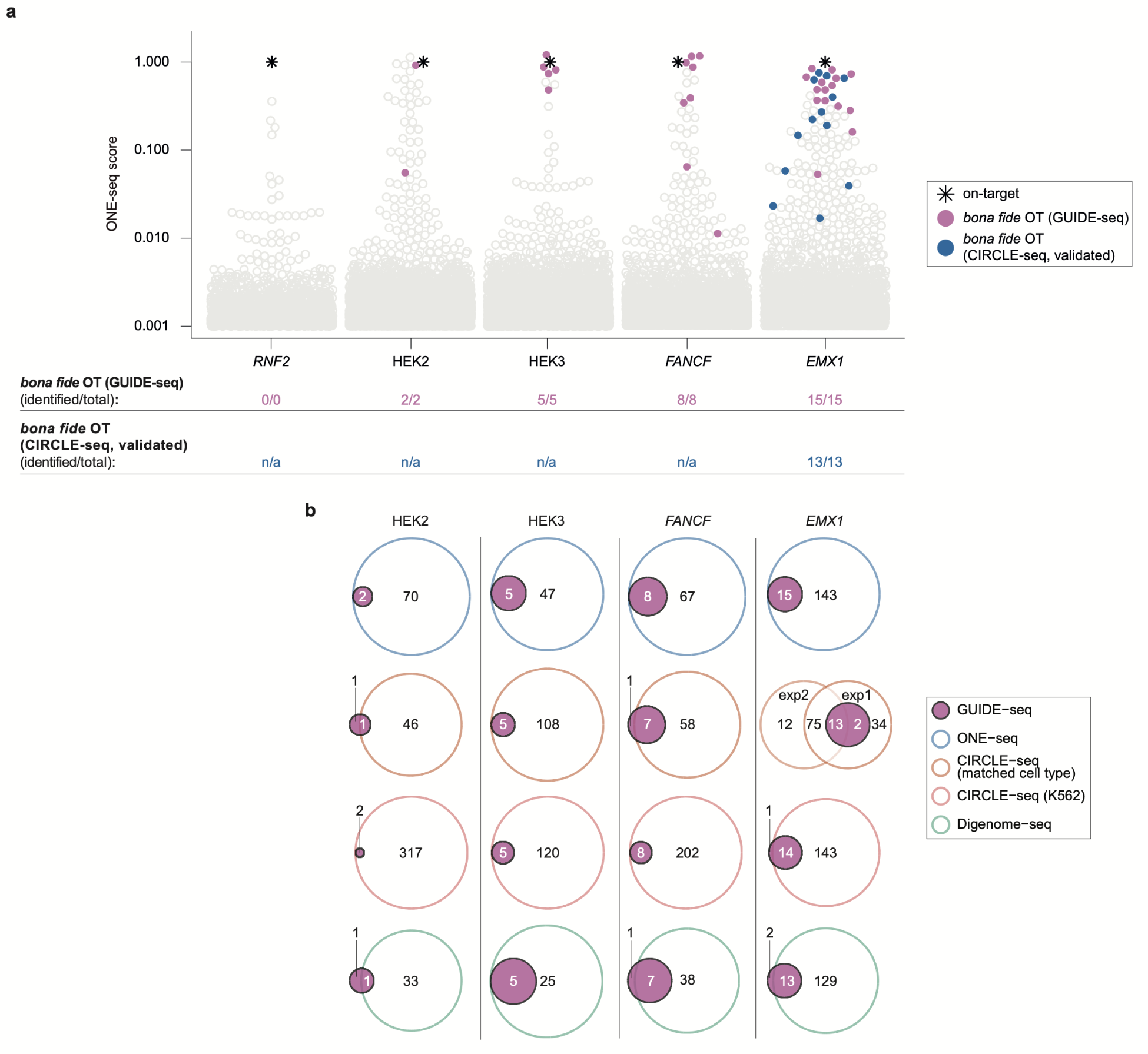
Results of ONE-seq off-target analysis with SpCas9 nucleases. **a,** Swarm plots showing ONE-seq nuclease scores for five previously analyzed SpCas9 gRNAs. Each circle represents an individual ONE-seq library member. Colored circles represent previously confirmed *bona fide* off-target sites. Sites with ONE-seq nuclease scores below 0.001 are not shown. n/a, no validation performed in previously published CIRCLE-seq study. **b,** Venn diagrams comparing abilities of ONE-seq, CIRCLE-seq, and Digenome-seq (open colored circles) to nominate *bona fide* off-target sites previously validated by GUIDE-seq (solid purple circles). All sites considered as validated by ONE-seq had ONE-seq nuclease scores >0.01.

**Extended Data Fig. 3.**
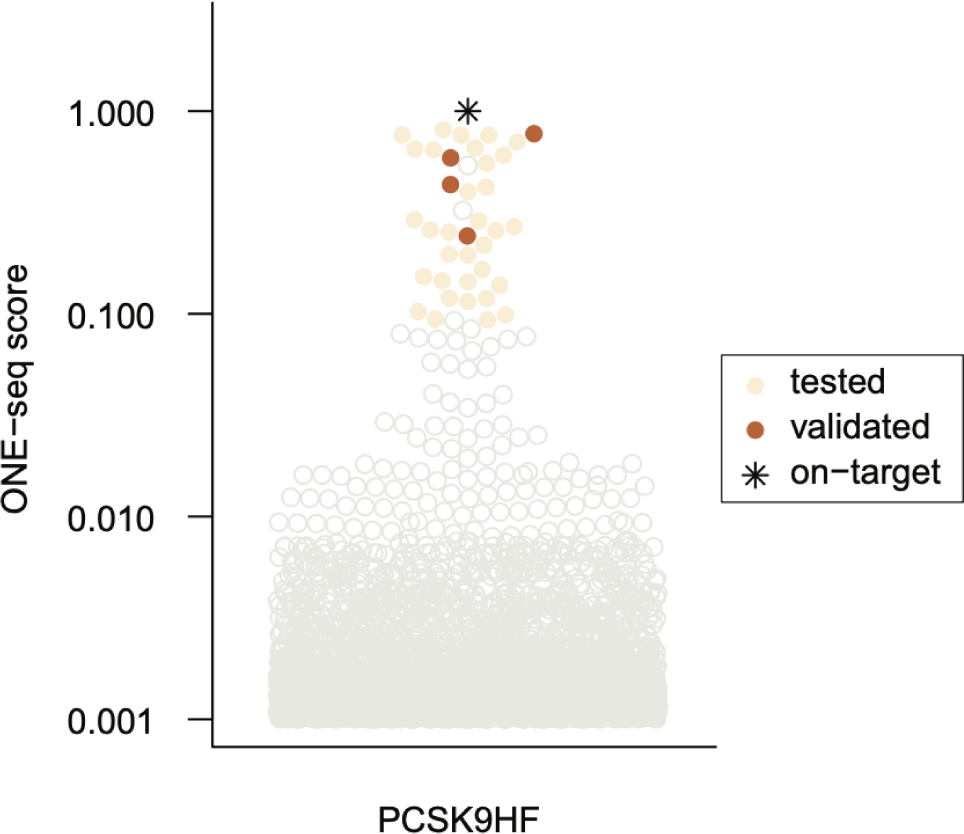
ONE-seq selections with SpCas9 and the PCSK9HF gRNA. Swarm plot showing ONE-seq nuclease scores for the SpCas9 PCSK9HF gRNA. Each circle represents an individual ONE-seq library member. Yellow circles represent off-target candidate sites that were tested by targeted amplicon sequencing from human hepatocyte DNA from chimeric mice. Orange circles represent sites tested by targeted amplicon sequencing that were validated as *bona fide* off-target sites in this study. The ONE-seq scores shown represent the average of four replicate ONE-seq experiments. Sites with ONE-seq nuclease scores below 0.001 are not shown.

**Extended Data Fig. 4.**
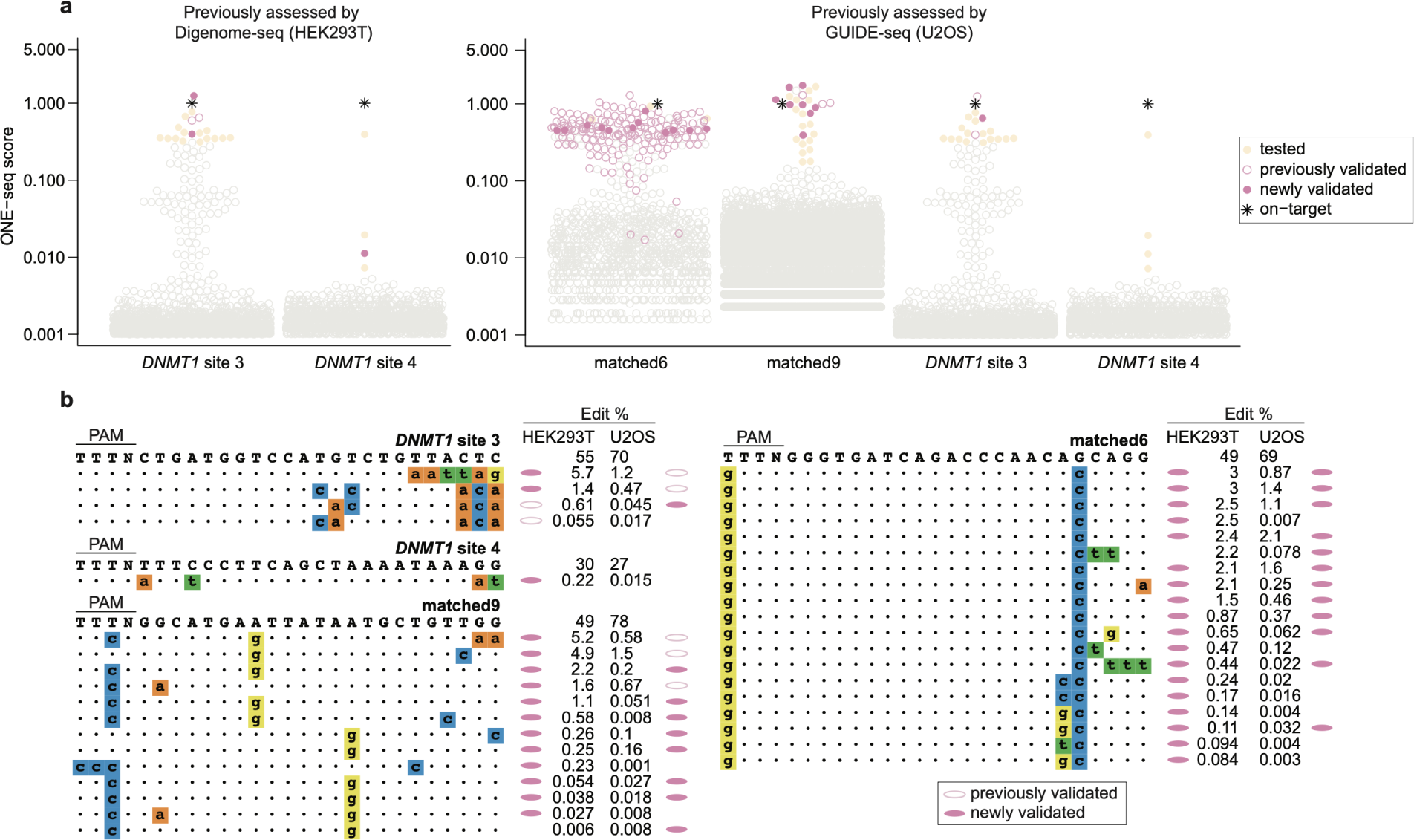
ONE-seq outperforms existing methods for nominating *bona fide* Cas12a nuclease off-targets in human cells. **a,** Swarm plots showing ONE-seq nuclease scores for Cas12a gRNAs previously characterized by Digenome-seq in HEK293T cells (left) and by GUIDE-seq in U2OS cells (right). Each circle represents an individual ONE-seq library member. Open pink circles and closed pink circles represent previously validated and newly validated Cas12a off-target sites, respectively. Other sites tested in this study by targeted amplicon sequencing are represented as closed yellow circles. Sites with ONE-seq nuclease scores below 0.001 are not shown. ONE-seq scores for matched9, DNMT3 and DNMT4 represent the average of duplicate ONE-seq experiments. **b,** Cas12a off-targets nominated by ONE-seq and tested and/or validated in HEK293T cells and/or U2OS cells are shown. Nucleotide sequences in bold at the top represent on-target sequences. Off-target sites assessed in cells are shown below the on-target site. Lower case nucleotides in colored boxes represent mismatches to the on-target site. Validation status of off-targets is shown by colored ovals. “Edit %” refers to the mean editing frequency from three independent replicates. 166 previously identified off-target sites for matched6 are not shown.

**Extended Data Fig. 5.**
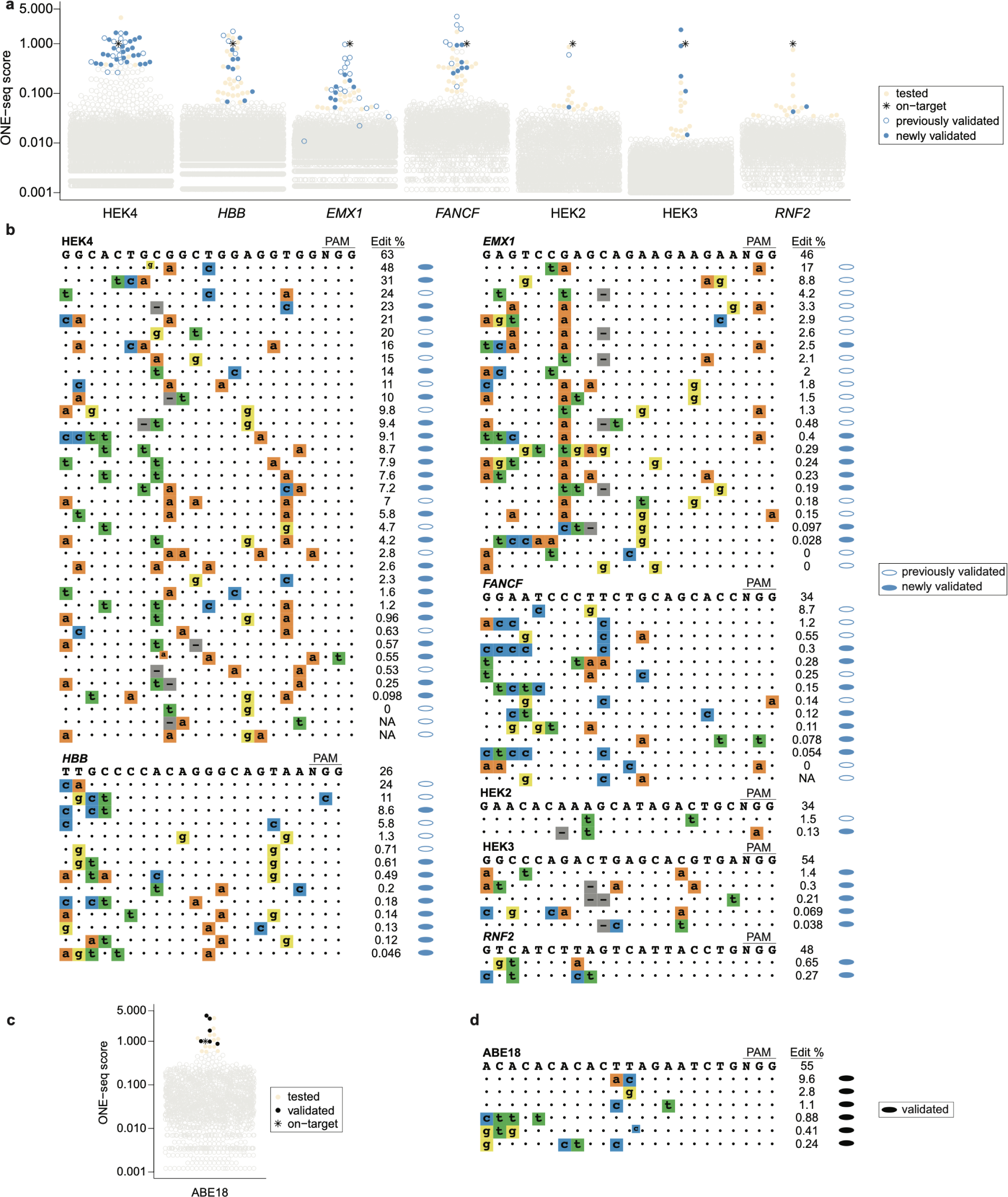
ONE-seq outperforms existing methods for nominating *bona fide* CBE BE3 off-targets in human cells. **a,** Swarm plots showing ONE-seq scores for the CBE BE3 with seven different gRNAs previously characterized by Di genome-seq. Each circle represents an individual ONE-seq library member. Sites with ONE-seq nuclease scores below 0.001 are not shown. **b,** BE3 off-targets nominated by ONE-seq for the seven gRNAs in **a** and validated in human HEK293T cells are shown. Nucleotide sequences in bold at the top represent on-target sequences. Off-target sites assessed in cells are shown below the on-target site. Lower case nucleotides in colored boxes represent mismatches to the on-target site. Validation status of off-targets is shown by colored ovals. “Edit %” refers to the mean editing frequency from three independent replicates. **c,** Swarm plots showing ONE-seq scores for the CBE BE3 with the ABE18 gRNA that was not previously characterized by Digenome-seq. Each circle represents an individual ONE-seq library member. Sites with ONE-seq nuclease scores below 0.001 are not shown. **d,** BE3 off-targets for the ABE18 gRNA nominated by ONE-seq and validated in human HEK293T cells are shown. Nucleotide sequences in bold at the top represent on-target sequences. Off-target sites assessed in cells are shown below the on-target site. Lower case nucleotides in colored boxes represent mismatches to the on-target site. Validation status of off-targets is shown by colored ovals. “Edit %” refers to the mean editing frequency from three independent replicates.

**Extended Data Fig. 6.**
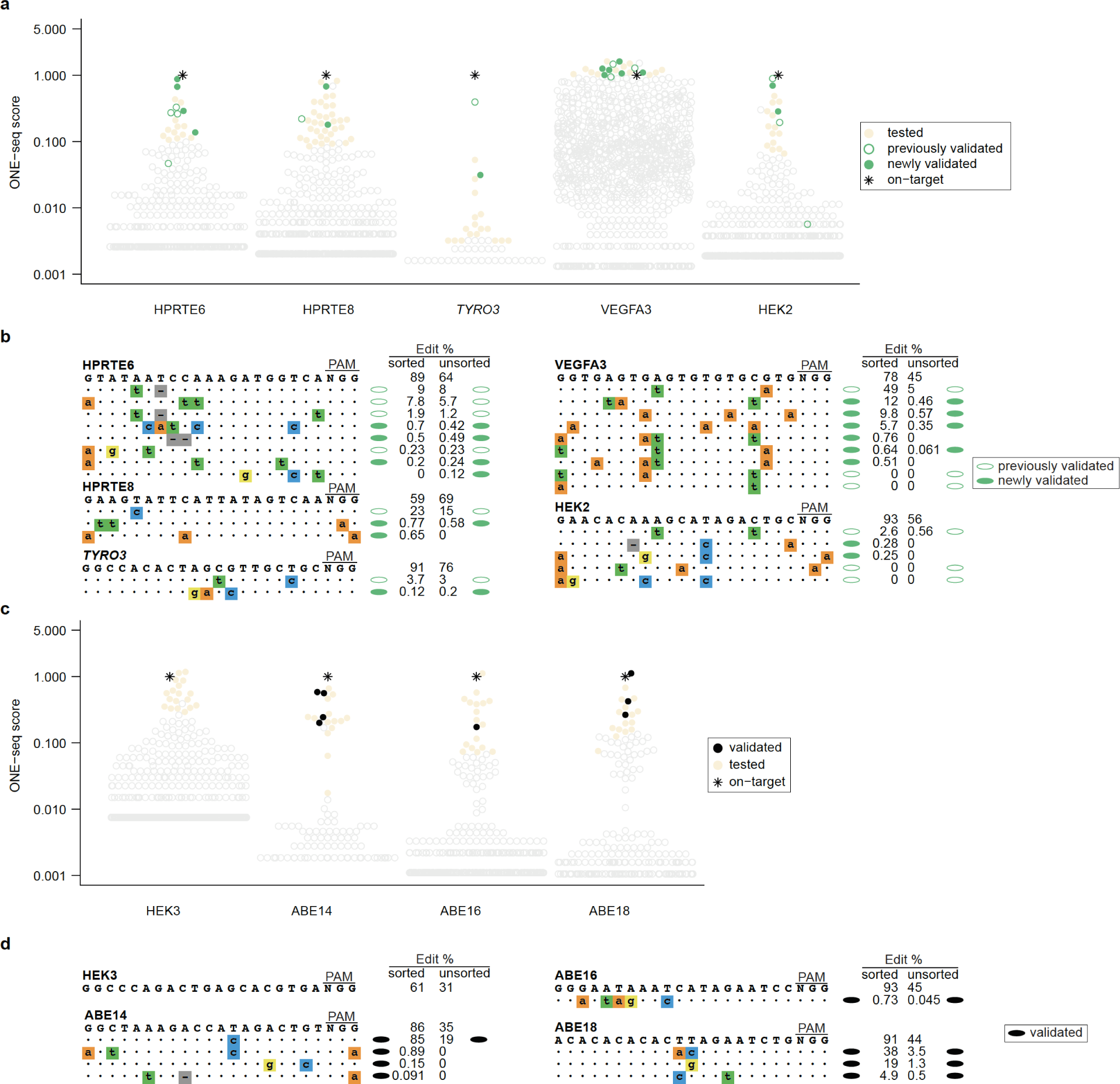
ONE-seq outperforms existing methods for nominating *bona fide* ABE 7.10 off-targets in human cells. **a,** Swarm plots showing ONE-seq scores for ABE 7.10 with five different gRNAs previously characterized by Digenome-seq or EndoV-seq. Each circle represents an individual ONE-seq library member. Sites with ONE-seq nuclease scores below 0.001 are not shown. **b,** ABE 7.10 off-targets nominated by ONE-seq for the five gRNAs in **a** and validated in human HEK293T cells are shown. Nucleotide sequences in bold at the top represent on-target sequences. Off-target sites assessed in cells are shown below the on-target site. Lower case nucleotides in colored boxes represent mismatches to the on-target site. Validation status of off-targets is shown by colored ovals. “Edit %” refers to the mean editing frequency from three independent replicates. **c,** Swarm plots showing ONE-seq scores for ABE 7.10 with four different gRNAs not previously characterized by Digenome-seq. Each circle represents an individual ONE-seq library member. Sites with ONE-seq nuclease scores below 0.001 are not shown. **d,** ABE 7.10 off-targets nominated by ONE-seq for the four gRNAs in **c** and validated in human HEK293T cells are shown. Nucleotide sequences in bold at the top represent on-target sequences. Off-target sites assessed in cells are shown below the on-target site. Lower case nucleotides in colored boxes represent mismatches to the on-target site. Validation status of off-targets is shown by colored ovals. “Edit %” refers to the mean editing frequency from three independent replicates.

**Extended Data Fig. 7.**
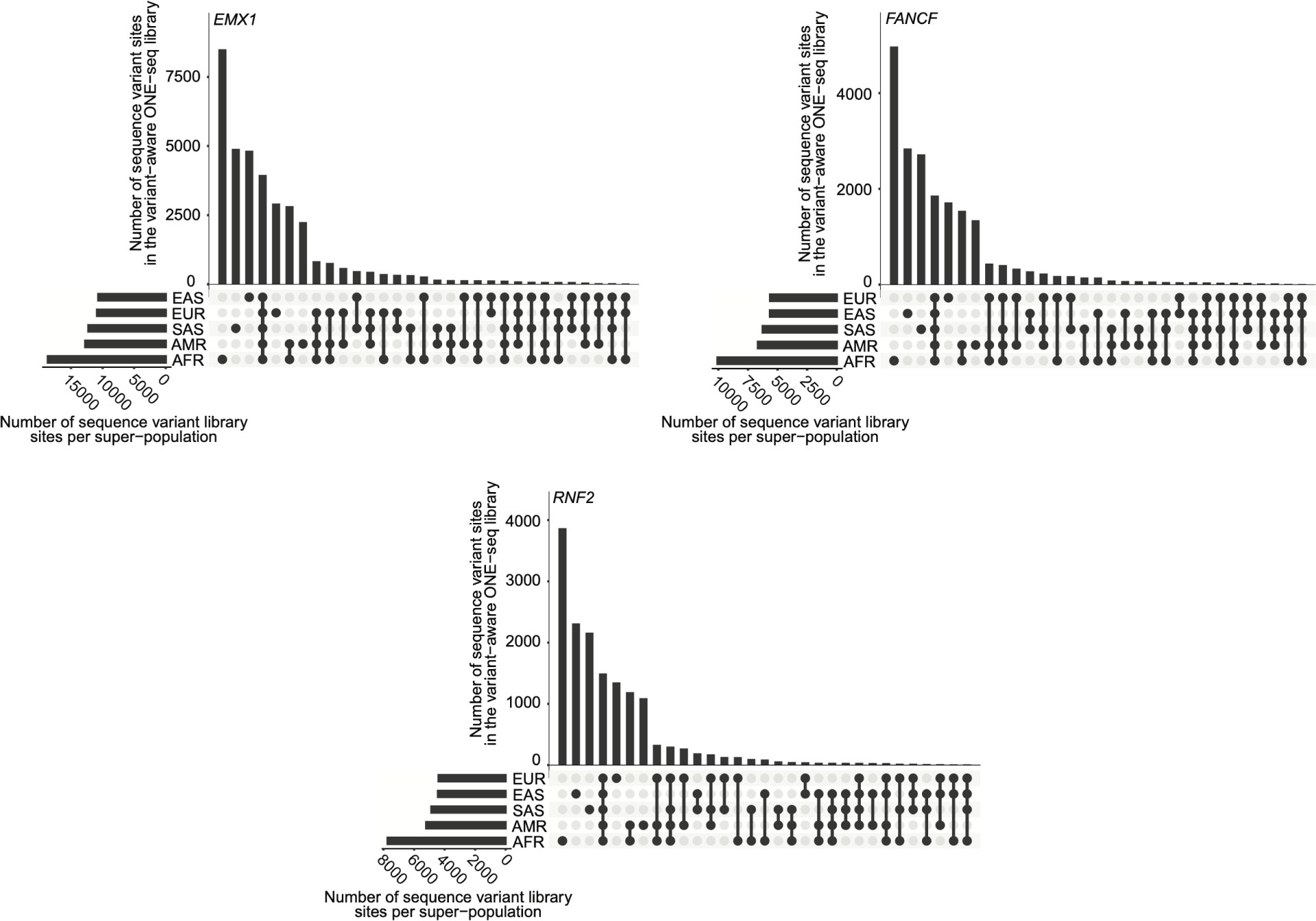
Super-population distribution of genomic sequence variants included in variant-aware ONE-seq libraries. Upset plots showing the distribution of genomic variants within super-populations for three variant-aware ONE-seq libraries. Each bar represents the count of genomic variants that are found in the super-populations indicated by black circles. The bar graphs to the bottom left represent the total number of genomic variants from the 1000 Genomes Project found in the European (EUR), East Asian (EAS), South Asian (SAS), American (AMR), and African (AFR) super populations, ordered from top to bottom by increasing frequency.

**Extended Data Fig. 8.**
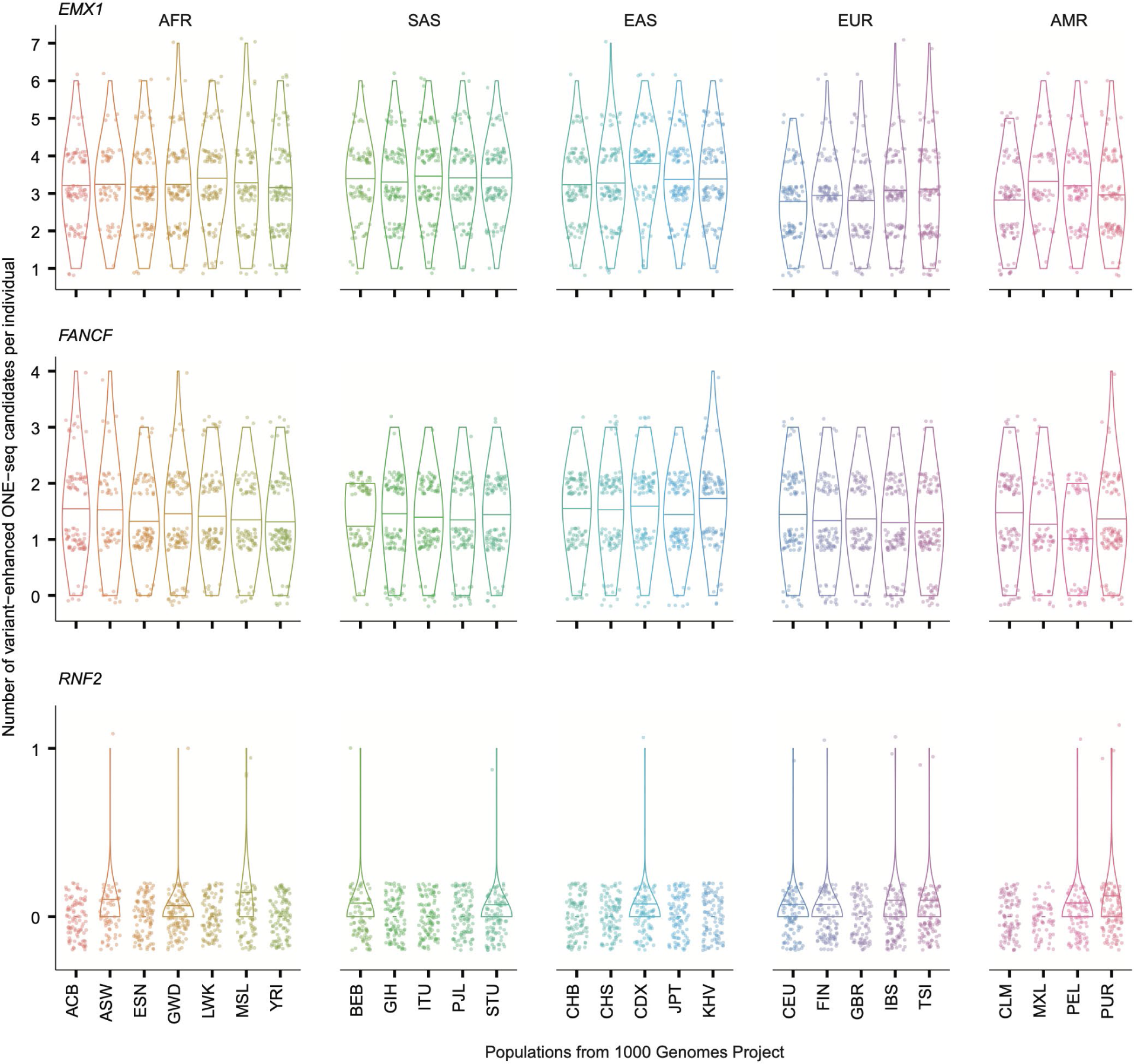
Numbers of variant-enhanced off-target candidates per individual stratified by super-population. Each dot represents an individual from the 1000 Genomes Project. The number of variant-enhanced off-target candidates per individual is shown, stratified by population. Three letter population abbreviations from the 1000 Genomes Project are shown for each population.

**Extended Data Fig. 9.**
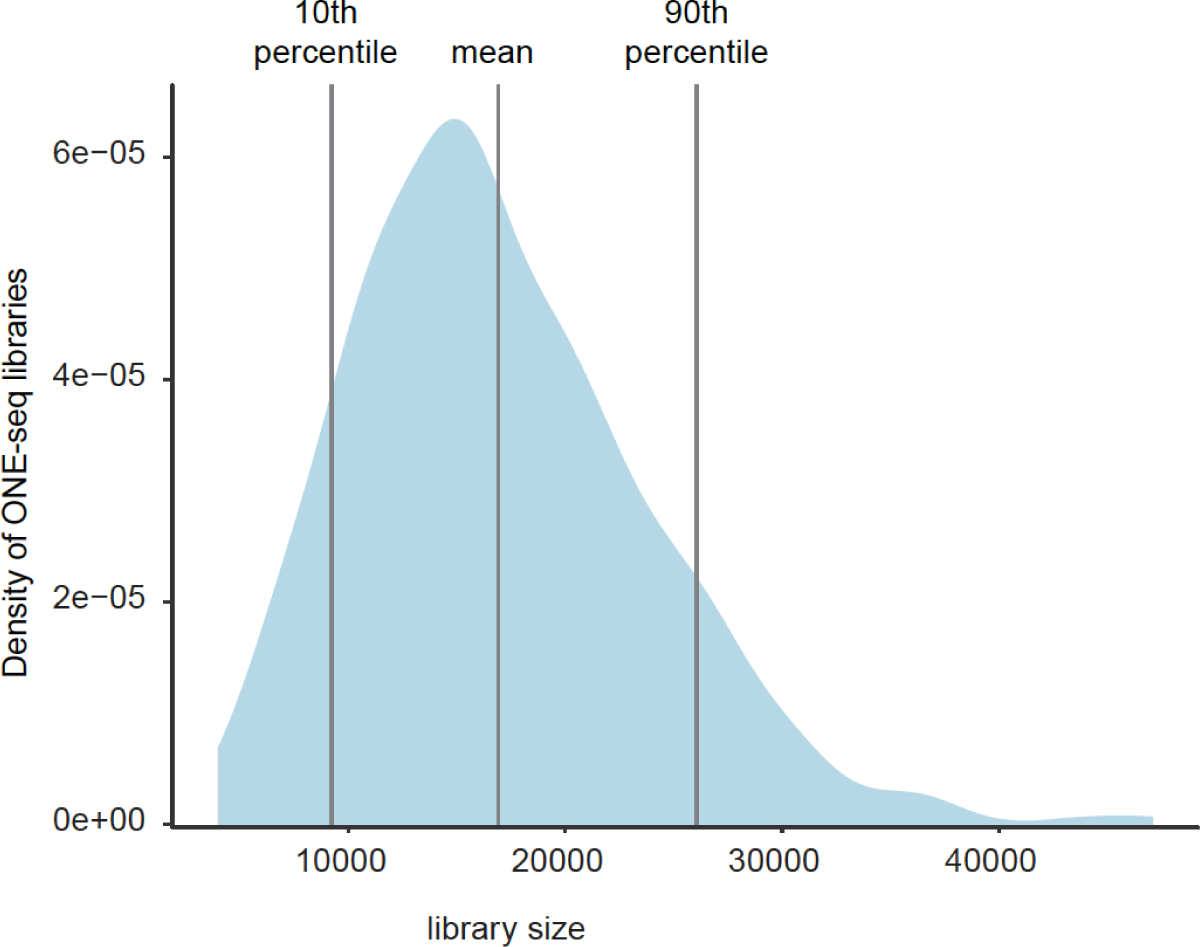
ONE-seq library sizes for 481 SpCas9 gRNAs. Distribution of ONE-seq library sizes for 481 gRNAs targeting 25 therapeutically relevant human genes. The graph reports the density of all ONE-seq libraries analyzed (vertical axis) with a given library size (horizontal axis).

