## Supplementary Information for "Global-scale CRISPR gene editor specificity profiling by ONE-seq identifies population-specific, variant off-target effects"

**Table of Contents**

**Supplementary Notes**

**Supplementary Result 1.** Comparison of SpCas9 gRNA target site coverage by CIRCLE-seq and ONE-seq libraries

**Supplementary Result 2.** Unsuccessful introduction of a DNA end blunting step into the CIRCLE-seq protocol

**Supplementary Algorithm.** ONE-seq analysis.

**Supplementary Tables**

**Supplementary Table 1.** Sequences of gRNAs used in this study.

**Supplementary Table 2.** Complexity of oligonucleotide libraries.

**Supplementary Table 3.** Uniformity of ONE-seq, CIRCLE-seq and WGS sequencing libraries.

**Supplementary Table 4.** PCSK9HF ONE-seq scores.

**Supplementary Table 5.** Composition of variant ONE-seq libraries.

**Supplementary Table 6.** Number of variant-sensitive ONE-seq off-targets.

**Supplementary Table 7.** Used oligonucleotides.

**Supplementary Table 8.** Sequences of used plasmids.

**Supplementary Table 9.** Sequence structure of ONE-seq libraries.

**Supplementary Notes**

**Supplementary Result 1.** Comparison of SpCas9 gRNA target site coverage by CIRCLE-seq and ONE-seq libraries

Following synthesis of ONE-seq libraries, we performed low-cycle amplification and sequencing of the pre-selection ONE-seq libraries to assess drop-out and uniformity of the chip-based oligonucleotide synthesis (**Extended Data Fig. 1; Supplementary Table 3**). To compare to CIRCLE-seq, we assessed the pre-selection composition of CIRCLE-seq libraries by targeting the common CIRCLE-seq linker with Cas9-gRNA ribonucleoprotein. Specifically, after circularization of genomic DNA^1^, we digested the CIRCLE-seq library with a gRNA/Cas9 complex targeted against a constant sequence present in circularized library members. We then ligated sequencing adapters and performed deep NGS with an Illumina NovaSeq. In the subsequent processing we only considered reads that contained the cleaved fragments of the CIRCLE-seq adapter on both paired reads. After this filtering step, we aligned the adapter-trimmed sequences to the reference hg19 human genome sequence.

To directly compare CIRCLE-seq and ONE-seq pre-selection library quality, we assessed uniformity and dropout of the genomic sequences that were defined in the ONE-seq libraries. We found that CIRCLE-seq libraries overall had a lower 90/10 ratio (2.8 +/- 0.4) and a higher dropout percentage (0.59% +/- 0.12%). Since dropout could be related to circularization, we also analyzed the same set of ONE-seq sites in standard whole genome sequencing preparations with and without PCR amplification. For both WGS sequencing methods, the 90/10 ratio was 2 +/- 0.2. The dropout percentage was 0.51% +/- 0.11% and 0.53% +/- 0.11% for the PCR and non-PCR library preparation, respectively. These results indicate that underrepresentation of ONE-seq sites in CIRCLE-seq libraries is most likely caused by intrinsic properties of the analyzed DNA rather than biases arising from circularization.

In addition to improved representation of sequences that are homologous to the on-target site in ONE-seq, it is notable that quality control of ONE-seq libraries can be done with a smaller-scale benchtop NGS machine. In contrast, because pre-selection CIRCLE-seq libraries contain unselected genomic DNA, pre-selection quality control requires access to WGS capabilities. In our experiments, two orders of magnitude fewer sequencing reads were needed to characterize pre-editing ONE-seq libraries compared to CIRCLE-seq (**data not shown**).

**Supplementary Result 2.** Unsuccessful introduction of a DNA end blunting step into the CIRCLE-seq protocol

Sequence-specific nucleases (such as ZFNs, TALENs, Cas9, and Cas12a) can leave blunt ends, staggered ends, or a mixture of both. Cas9 nuclease mostly leaves blunt DNA ends but has been reported to introduce single base pair overhangs at certain target sites ^2, 3^. Other designer nuclease types such as Cas12a or homing endonucleases, such as I-PpoI or I-SceI, generally leave staggered DNA ends after cleavage. Therefore, we tried to adapt the CIRCLE-seq protocol to introduce a blunting step. Following circularization of DNA according to the CIRCLE-seq protocol, we introduced a blunting step directly after Cas9 cleavage. When sequencing the blunted CIRCLE-seq libraries ~10-fold fewer reads passed the CIRCLE-seq pipeline in comparison to a non-blunted CIRCLE-seq reaction. Although we do not know why the extra blunting step reduces the number of useable reads so dramatically, we hypothesize that this 90% decrease in useable reads results from conversion of double-nicked circles present in the background of the selections, unrelated to Cas9 activity, into a ligation-compatible form through the additional blunting step.

**Supplementary Algorithm.** ONE-seq analysis.

1) Quality filtering, adapter trimming and merging of sequencing reads.
a) Trim reads using trimmomatic version 0.36^4^ in paired end mode with the following parameters: ‘java -jar trimmomatic-0.36.jar PE -phred33 -trimlog {trimlog path} -basein {inputfile path} -baseout {output file path} ILLUMINACLIP:{adapter file path}:2:30:10:1:true LEADING:0 TRAILING:0 SLIDINGWINDOW:4:30 MINLEN:36 2>&1 | tee {output path}’ .
b) Merge reads with flash version 1.2.11^5^ with the following parameters: ‘flash --max-mismatch-density=0.25 –output-directory={output path} –output-prefix=merged_{suffix} --compress --max-overlap=160 ’ {input file fw path} {input file rev path} 2>&1 | tee {log path}_flashlog.txt.
2) Annotation of sequencing reads.
a) Check the beginning of R1 for the presence of the ONE-seq adapter and assign a boolean value to the linkerbool attribute of the sequencing read object depending on if the adapter is found or not.
b) Check the beginning of R2 for the presence of the expected primer sequence and assign a boolean value to the primerbool attribute of the sequencing read object depending on if the primer sequence is found or not.
c) Search for constant1 and constant2 (proto side amplification) or constant5 and constant6 (PAM side amplification) in the sequencing read object (**Supplementary Table 11)** and determine the length of the sequence in between the two constant sequences. If the sequence is the expected length of the UMI (default: 11bp) assign the UMI attribute of the sequencing read object to the intervening sequence otherwise set the UMI attribute to ‘None’.
d) Search for constant2 and constant3 (proto side amplification) or constant4 and constant5 (PAM side amplification) in the sequencing read object (**Supplementary Table 11**) and determine the length of the sequence in between the two constant sequences. If the sequence is the expected length of the ONE-seq library barcode (default: 14bp), set the barcode attribute of the sequencing read object to the intervening sequence, otherwise set the barcode attribute to ‘None’.
e) Search for constant 3 and the ONE-seq adapter sequence (proto side amplification) or for contant4 and the ONE-seq adapter sequence (PAM side amplification) in the sequencing read object. If both sequences are found assign the sequence in between the two constant sequences as the target attribute of the sequencing read object, otherwise set the target attribute to ‘None’.
3) Filter out sequencing read objects that fulfill one of the following criteria.
a) Seq_object.linkerbool != True. (ONE-seq adapter not found)
b) Seq_object.primerbool != True. (Amplification primer not found)
c) Seq_object.UMI == None. (UMI couldn’t be assigned)
d) Seq_object.barcode == None. (ONE-seq barcode couldn’t be assigned)
e) Seq_object.target == None. (target couldn’t be assigned)
4) Load sequencing read objects into ONE-seq library.
a) Generate library member objects from the ONE-seq library file.
b) Compare barcodes of library member objects with barcodes of sequencing read objects and append sequencing read objects to the foundseqs attribute of library member objects if the sequencing read and library member barcodes are identical.
5) Identify sequencing read objects with expected *in vitro* editing outcomes.
a) Calculate the alignment position of the ONE-seq adapter for each sequencing read object by comparing the target attribute of the sequencing read object with the full oligonucleotide sequence of the library member object to which the sequencing read object was assigned.
b) Check if the alignment position of the ONE-seq adapter is compatible with an expected in vitro editing outcome. This step varies depending on the editor.

Cas9:

- Declare a correct ONE-seq adapter alignment position if the ONE-seq sequencing adapter ligated within a two-sided 2bp window around the expected cleavage site of Cas9 (for protospacer and PAM side amplification).

Cas12:

- Declare a correct ONE-seq adapter alignment position if the ONE-seq sequencing adapter ligated within a two-sided 2-3 bp window around the expected cleavage site of Cas12 (for protospacer and PAM side amplification).

CBE:

- Declare a correct ONE-seq adapter alignment position if the ONE-seq adapter ligated within a two-sided 2 bp window around the nicking site (for the protospacer side amplification).
- Declare a correct alignment if the ONE-seq adapter ligated at a position following a C in the oligonucleotide sequence (for the PAM side amplification).

ABE:

- Declare a correct ONE-seq adapter alignment position if the ONE-seq adapter ligated within a two-sided 2 bp window around the nicking site and if an A -> G edit is detected in the target sequence (for the protospacer side amplification)

Note: This approach is enabled by the properties of Endonuclease V, which cleaves the second phosphodiester bond 3’ of an inosine, thus conserving the A -> G edit in the sequenced DNA fragment.

6) Calculate ONE-seq scores.

- Count the number of sequencing reads with correct ONE-seq adapter alignment positions for each ONE-seq library member.
- Calculate the sided ONE-seq score by dividing the sequencing read count (with correct adapter ligation position) of a ONE-seq library member with the sequencing read count (with correct adapter ligation position) of the ONE-seq library member that represents the on-target sequence of a specific gene editor.
- Calculated the combined ONE-seq score by averaging the sided protospacer-side and PAM-side ONE-seq score.
- Note: For ABE, only the protospacer-side ONE-seq score is considered, since the editing outcome (A to G change) is directly visible in the protospacer-side amplification.

**Supplementary Information References**
