## Supplementary Figure 1 for "Global-scale CRISPR gene editor specificity profiling by ONE-seq identifies population-specific, variant off-target effects"

FACS raw data and gating examples for different experimental conditions.

The following figures depict the sorting strategy for control and base editor expressing HEK293T cells and control and Cas9 expressing lymphoblastoid cells. The image data was generated at the MGH Molecular Pathology Flow Cytometry Core Facility on a BD FACSAria II or Fusion cytometer using BD FACSDiva v. 6.1.3 and BD FACSDiva 8.01. The data shown is taken from original batch analysis files. We performed sorting for GFP for HEK293T cells and GFP and mCherry sorting for lymphoblastoid cells. The sorting strategy involved doublet exclusion, gating on the population of interest and sorting for the respective fluorochrome. For GFP sorting the x-axis shows FITC which closely matches the spectral profile of GFP. For mCherry sorting the x-axis shows PE-Texas Red, which closely match the spectral characteristics of mCherry.

### LCL GFP mCherry

#### LCL Negative Control (no transfection)

Doublet exclusion

Gating on population

Gating for GFP+/mCherry+ cells

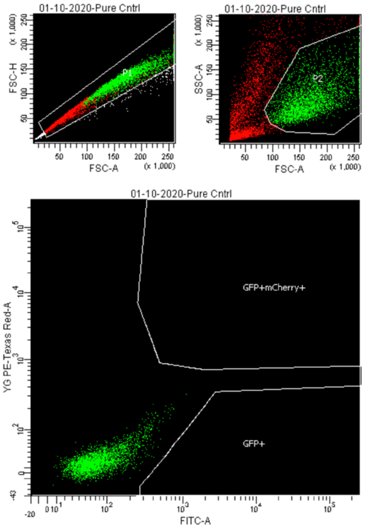


Tube: Pure Cntrl

| Population | #Events | %Parent | %Total |
|---|---|---|---|
| All Events | 10,000 | #### | 100.0 |
| P1 | 6,374 | 63.7 | 63.7 |
| P2 | 3,062 | 48.0 | 30.6 |
| GFP+ | 2 | 0.1 | 0.0 |
| GFP+mCherry+ | 0 | 0.0 | 0.0 |

Population statistics

---

#### LCL All GFP mCherry
#### Co-transfection with Cas9-P2A GFP and mCherry-sgRNA

Doublet exclusion

Gating on population

Gating for GFP+/mCherry+ cells

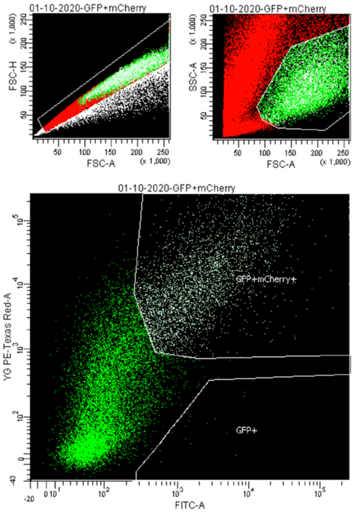


Tube: GFP+mCherry

| Population | #Events | %Parent | %Total |
|---|---|---|---|
| All Events | 100,000 | #### | 100.0 |
| P1 | 61,036 | 61.0 | 61.0 |
| P2 | 16,555 | 27.1 | 16.6 |
| GFP+ | 16 | 0.1 | 0.0 |
| GFP+mCherry+ | 3,716 | 22.4 | 3.7 |

Population statistics

### HEK293T GFP

#### HEK293T Negative Control (no transfection)

Doublet exclusion

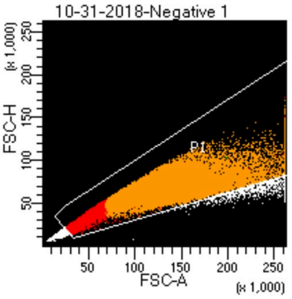


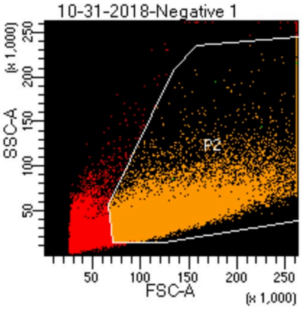


Gating on population

Gating for GFP+ cells

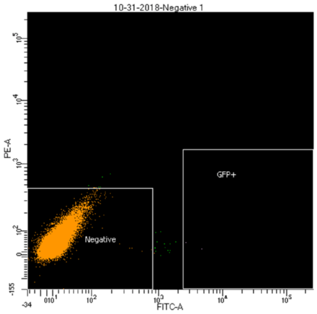


BD FACSDiva 8.0.1

| | |
|---|---|
| Experiment Name: | Experiment_003 |
| Specimen Name: | 10-31-2018 |
| Tube Name: | Negative 1 |
| Record Date: | Oct 31, 2018 2:45:26 PM |
| $OP: | JoungLab |
| GUID: | 8087c930-1690-43f3-8e0f-1d5d... |

| Population | #Events | %Parent | FITC-A Geo Mean |
|---|---|---|---|
| GFP+ | 2 | 0.0 | 3,472 |
| Negative | 67,984 | 100.0 | #### |
| Neg | #### | #### | #### |
| P3 | #### | #### | #### |

Tube: Negative 1

| Population | #Events | %Parent | %Total |
|---|---|---|---|
| All Events | 115,504 | #### | 100.0 |
| P1 | 103,787 | 89.9 | 89.9 |
| P2 | 68,005 | 65.5 | 58.9 |
| GFP+ | 2 | 0.0 | 0.0 |
| Negative | 67,984 | 100.0 | 58.9 |
| Neg | #### | #### | #### |
| P3 | #### | #### | #### |

Population statistics

---

#### HEK293T top ~25% total GFP
#### Transfection with ABEmax-P2A GFP

Doublet exclusion

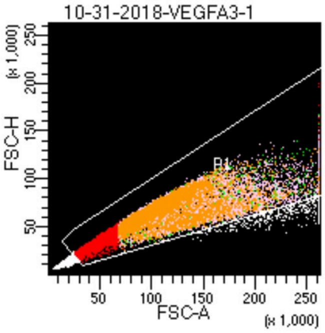


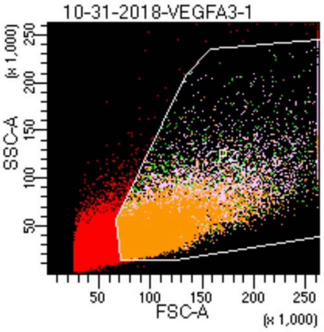


Gating on population

Gating for GFP+ cells

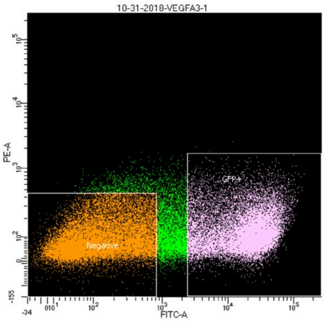


BD FACSDiva 8.0.1

| | |
|---|---|
| Experiment Name: | Experiment_003 |
| Specimen Name: | 10-31-2018 |
| Tube Name: | VEGFA3-1 |
| Record Date: | Oct 31, 2018 2:54:52 PM |
| $OP: | JoungLab |
| GUID: | dca404fe-4071-4962-adfb-6b22... |

| Population | #Events | %Parent | FITC-A Geo Mean |
|---|---|---|---|
| GFP+ | 21,644 | 41.9 | 16,510 |
| Negative | 24,425 | 47.3 | #### |
| Neg | #### | #### | #### |
| P3 | #### | #### | #### |

Tube: VEGFA3-1

| Population | #Events | %Parent | %Total |
|---|---|---|---|
| All Events | 100,000 | #### | 100.0 |
| P1 | 87,792 | 87.8 | 87.8 |
| P2 | 51,603 | 58.8 | 51.6 |
| GFP+ | 21,644 | 41.9 | 21.6 |
| Negative | 24,425 | 47.3 | 24.4 |
| Neg | #### | #### | #### |
| P3 | #### | #### | #### |

Population statistics
